# Reversible and Causal Epigenetic Information Loss in Liver Aging and Disease

**DOI:** 10.1101/2025.02.24.639802

**Authors:** Roni B. Shtark, Naor Sagy, Noga Korenfeld, Maayan Gal, Ido Goldstein, Daniel Z. Bar

**Affiliations:** Department of Oral Biology, Goldschleger School of Dental Medicine, The Faculty of Medical and Health Sciences, Tel Aviv University, 69978 Tel Aviv, Israel; Institute of Biochemistry, Food Science and Nutrition. The Robert H. Smith Faculty of Agriculture, Food and Environment. The Hebrew University of Jerusalem. 229 Herzl Street, Rehovot 7610001, Israel

**Author notes:** Equal contribution (Roni B. Shtark, Naor Sagy).

## Abstract

The loss of epigenetic information has been proposed as a driver of aging and diseases, but the reversibility and causality of this process remain underexplored. Here we analyze liver-unique methylation sites - genomic loci that show distinct methylation patterns in the liver compared to other tissues. Upon disease progression, these sites overwhelmingly regress toward the pan-tissue average. In addition, we demonstrate that this regression also occurs in a majority of these sites during normal aging. Using Mendelian randomization analysis, we identify significant enrichment of liver-unique methylation sites in causal aging-associated loci, particularly sites that are highly methylated in healthy liver. Remarkably, repeated fasting, a metabolic intervention known to improve liver function, partially restores the liver-unique methylation patterns at these sites. This restoration also occurs in isolated hepatocytes subjected to fasting-mimicking conditions, suggesting the effect is cell-autonomous rather than due to changes in tissue composition. The liver-unique methylation sites are enriched for binding sites of key metabolic transcription factors and show significant overlap with genetic variants associated with liver disease risk, suggesting a mechanistic link between epigenetic information loss and liver dysfunction. Our findings establish epigenetic information loss as both a marker and mediator of liver aging and disease, while demonstrating its potential reversibility through metabolic interventions.

**Graphical abstract: Reversible information loss at liver-unique methylation sites:** Liver-unique sites, showing higher (UH) or lower (UL) methylation levels, regress to the pan-tissue average upon aging and disease. UH are enriched for methylation sites causal to the aging process, while UL are enriched for liver-specific enhancers and PPAR-α binding sites. Upon repeated fasting, both UL and UH diverge away from the pan-tissue average, partially restoring the more youthful and disease-free epigenetic state.

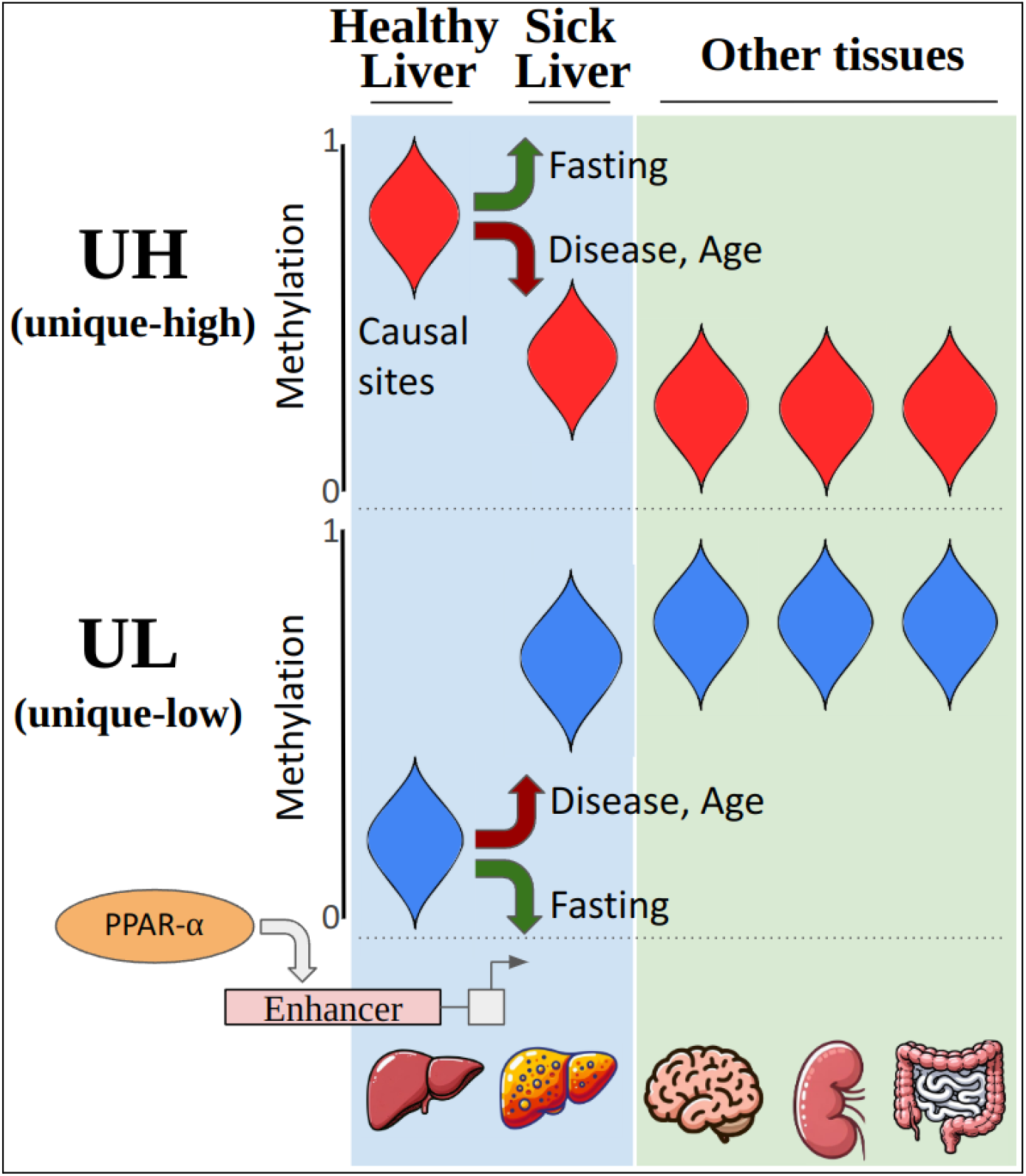

## Introduction

### Liver diseases and aging

The liver is a vital organ with remarkable regenerative capacity, yet it remains susceptible to age-related deterioration and disease. With advancing age, liver function generally declines, and aging is associated with increased severity and poor prognosis of multiple liver diseases ^1–3^. This decline manifests through various molecular and cellular changes, including alterations in hepatic blood flow, immune responses, and regenerative potential ^4^. Recent evidence suggests that the loss of epigenetic information—measured here via the regression of tissue-specific methylation patterns toward the pan-tissue average— accompanies aging and some diseases ^5^. However, whether this loss is reversible, and to what extent it causally contributes to liver aging and dysfunction, remains unknown. Understanding the reversibility and causality of epigenetic information loss could open new therapeutic avenues for age-related liver conditions.

### Epigenetic clocks and liver aging

DNA methylation patterns at specific loci change predictably with age, enabling the development of epigenetic clocks that measure chronological and biological age. These clocks, originally developed using blood samples, typically combine methylation levels at multiple CpG sites using elastic net regression or similar machine learning approaches ^6–8^. Recent advances using Mendelian randomization have identified specific methylation sites that causally contribute to aging, distinguishing between detrimental (DamAge) and adaptive (AdaptAge) age-related changes ^9^. Epigenetic clocks spanning multiple tissues, and even pan-tissue and pan-species have been developed ^10,11^. In mouse models, liver-specific methylation clocks have been constructed ^12^, though their relationship with liver function remains unclear. Understanding how these aging-associated methylation changes interact with tissue-specific epigenetic patterns could provide insights into the mechanisms of liver aging.

### Tissue-specific methylation and epigenetic information loss

The epigenetic theory of aging suggests that the erosion of methylation patterns contributes to aging ^13–15^. Recent evidence indicates that this loss of epigenetic information affects tissue function during aging and pathological conditions ^5,16–18^. Restoration of a youthful epigenetic state was demonstrated using exogenous factors ^17,19,20^. Multiple studies have identified tissue-specific and cell-type specific methylation patterns using combinations of various experimental and computational approaches ^21^. Here, we focus on a specific subset of methylation sites identified through a comparative tissue analysis approach ^5^. Due to differences in selection criteria, statistical thresholds, and reference tissue sets, these sites differ substantially from previously reported tissue and cell specific methylation sites. To avoid confusion with existing nomenclature, we refer to these as tissue-unique methylation sites. These sites show two distinct patterns: unique-low (UL) sites have lower methylation values in a specific tissue compared to other tissues, while unique-high (UH) sites show higher tissue-specific methylation. During aging and disease, these sites regress toward the average methylation levels found across tissues. This phenomenon has been observed across multiple organs and pathologies, including kidney disease ^18^. In the liver, these changes correlate with decreased organ function, though the causal relationship remains unexplored ^5^.

### Dietary interventions and Epigenetic Changes

Several dietary interventions, such as dietary/caloric restriction (CR) and intermittent fasting (IF), have shown promise in protecting against and ameliorating liver diseases, including inflammation and fibrosis. In both humans and animal models, these interventions have demonstrated the ability to delay or even prevent liver deterioration ^22^. Notably, CR and IF increase lifespan in multiple animal models ^23,24^. These dietary interventions have been shown to influence DNA methylation, histone acetylation, and chromatin remodeling, potentially delaying age-associated epigenetic changes ^25–27^. However, the exact relationship between these interventions and tissue-specific epigenetic modifications remains unclear ^28^.

### Study aims and findings

To explore the reversibility and causality of epigenetic information loss, we investigated tissue-unique methylation sites, where information loss can be easily observed (Graphical abstract). Our analysis revealed that liver-unique methylation sites are enriched for age-correlated sites and are found near disease-associated SNPs. Furthermore, these sites show significant overlap with detrimental age-related methylation sites identified through Mendelian randomization. Importantly, we demonstrate that metabolic interventions in mice can partially reverse these changes, both in whole livers and isolated hepatocytes, suggesting epigenetic information loss is at least partially reversible in the liver.

## Results

### Cirrhotic liver methylation patterns are enriched for liver-unique methylation sites

To validate the relationship between tissue-specific methylation patterns and liver disease, we analyzed methylation changes in an independent dataset of cirrhotic, hepatocellular carcinoma (HCC), and control human livers ^29^. Of 1,750 differentially methylated sites identified, 1,615 were specific to HCC samples. Of these HCC-specific sites, only a small fraction (42 sites; <3%) overlapped with liver-unique methylation sites. In contrast, among the 120 sites showing differential methylation in cirrhosis, we observed a striking enrichment of liver-unique methylation sites (53 sites; 44%, p = 0.0001). This enrichment was independent of HCC status and showed a clear directional pattern: hypomethylated sites in cirrhosis overlapped exclusively with UH sites, while hypermethylated sites overlapped exclusively with UL sites (**Fig. 1**; **Supp. Tables 1**,**2**). These findings demonstrate that liver cirrhosis is associated with regression of liver-unique methylation patterns toward the mean methylation levels observed across tissues, consistent with disease-associated epigenetic information loss.

**Figure 1:**
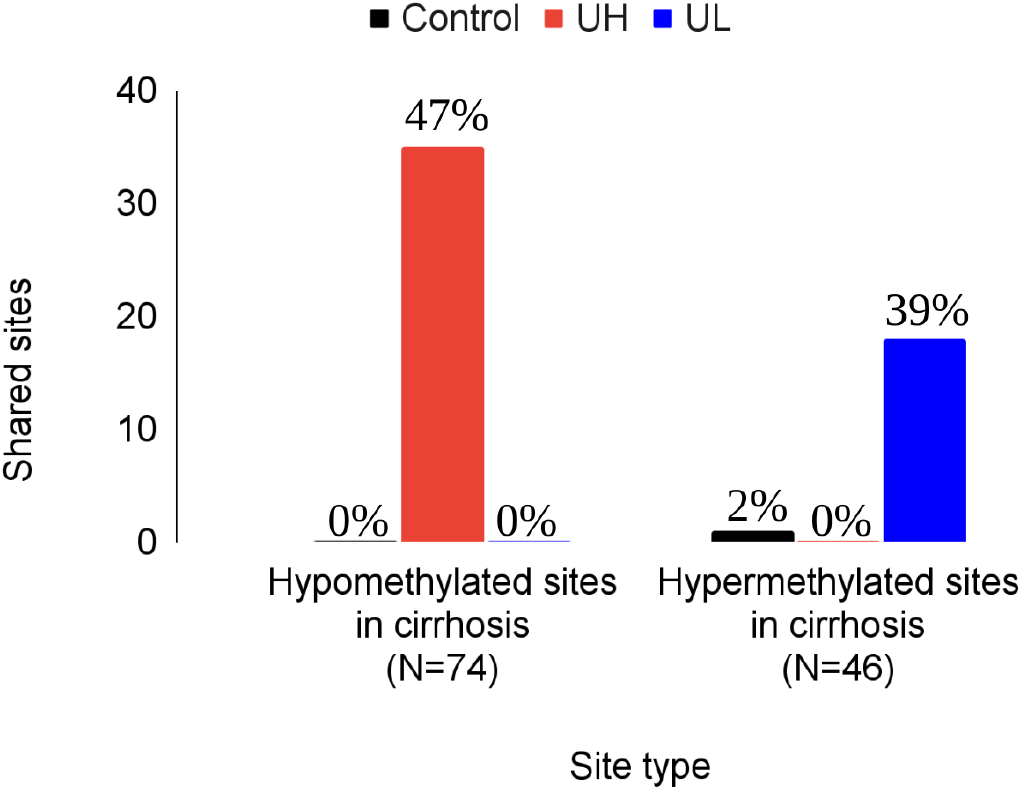
Liver-unique methylation sites are enriched in cirrhotic liver methylation patterns, which regress to the mean. Overlap of methylation sites hypomethylated (left) and hypermethylated (right) in cirrhosis with UH, UL, and control sites reveals a significant overlap of hypomethylated with UH, and hypermethylated with UL, suggesting a regression to the mean. A single control site overlapped with the hypomethylated sites. Monte Carlo simulation demonstrates that this observed enrichment of liver-unique methylation sites is statistically significant (p = 0.0001).

### Disease-associated genes are enriched for liver-unique methylation sites

To investigate potential causal relationships between tissue-specific epigenetic regulation and liver disease, we examined the spatial distribution of liver-unique methylation sites relative to genes containing established genetic risk variants for liver diseases ^30^. Among 17 genes previously identified as containing causal variants for various liver pathologies, we found significant enrichment of liver-unique methylation sites within 20 Kb of these loci. Specifically, 7 genes harbored a total of 13 liver-unique methylation sites (**Fig. 2A**). These genes encode proteins central to liver metabolism: PNPLA3 (hepatic stellate cell lipase), MTTP (VLDL triglyceride loading), ADH1B (alcohol metabolism), GCKR (glucokinase regulatory protein), MOSC1 (amidoxime reduction), Apo-E (fat metabolism), and GPAM (glycerolipid synthesis). The enrichment was highly significant compared to random control regions (p = 0.0001) and decreased with distance from the genetic variants (**Supp. Table 3**).

**Figure 2:**
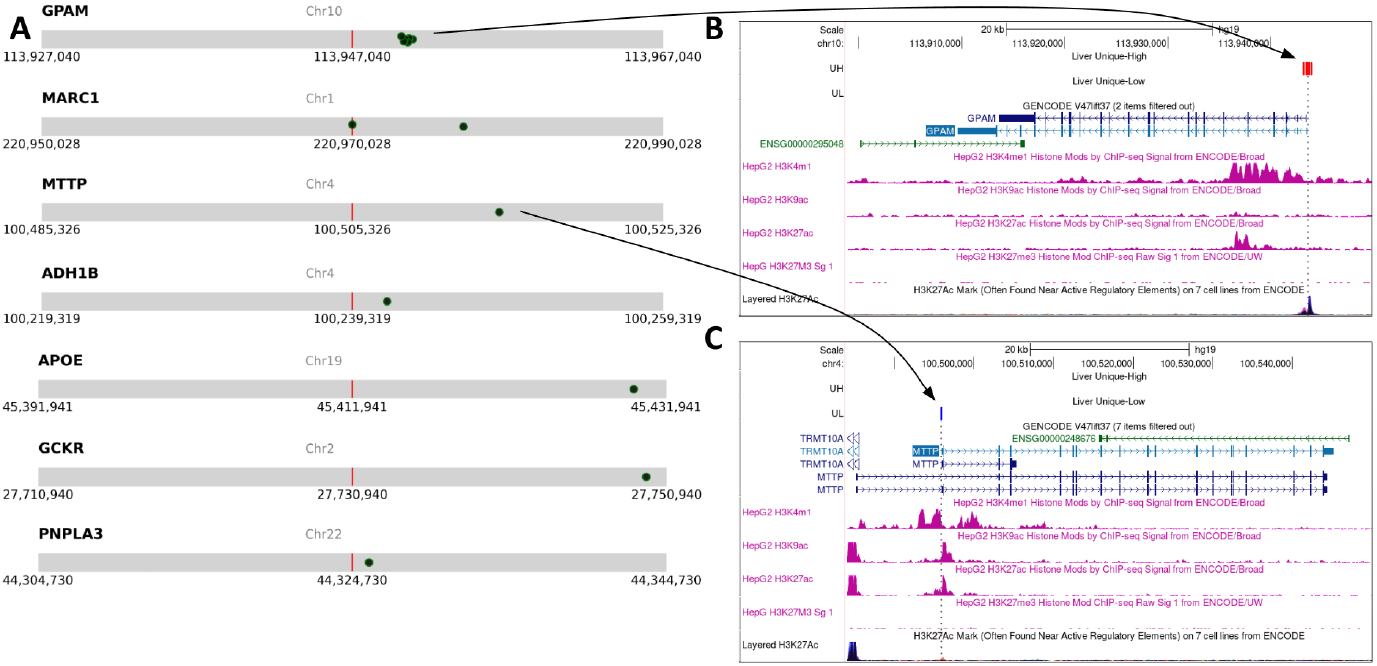
Liver-unique methylation sites are enriched in proximity to pathogenic SNPs. **(A)** Distribution of liver-unique methylation sites (black circle) relative to risk factor SNPs for liver diseases (red line). **(B)** Genome-browser view of the six UH methylation sites (red, indicated by an arrow), which overlap the TSS and first exon of GPAM, as well as H3K27ac in multiple cell lines. However, liver-derived HepG2 cells do not show H3K27ac at that location. **(C)** Genome-browser view of the UL site (blue, indicated by an arrow) adjacent to the TSS of MTTP, as well as enhancer-associated chromatin marks including H3K27ac and H3K4me1. The other 7 cell lines from ENCODE do not show significant H3K27ac levels at this location.

To understand the functional implications of these liver-unique methylation sites, we examined their relationship with gene expression and chromatin states. The presence of liver-unique methylation sites did not necessarily correspond to liver-specific expression. For instance, GPAM, despite having 6 UH sites near its transcription start site (TSS), is expressed in multiple tissues. Interestingly, while its TSS shows H3K27ac enrichment in several cell lines, this mark is notably absent in liver-derived HepG2 cells (**Fig. 2B**). Conversely, MTTP, which is expressed predominantly in liver and gastrointestinal tissue, contains a single UL site near its TSS that coincides with active chromatin marks (H3K27ac, H3K9ac, and H3K4me1) specifically in HepG2 cells but not in non-liver cell lines (**Fig. 2C**). These results suggest that causal genetic variants for liver disease often present near regions with liver-unique methylation patterns, though the mechanistic basis of this association requires further investigation.

### Characterization of liver-unique methylation sites in mice

Mouse experiments offer better control over genetic and environmental effects, as well as sample access. To expand our analysis, we generated a dataset of liver-unique methylation sites in mice, using methods similar to ^5,21^, based on public data from GSE184410 ^31,32^. We identified 6038 liver-unique methylation sites, of these 38% were UH and 62% were UL (**Supp. Table 4**). Sequence analysis using HOMER revealed distinct transcription factor motif enrichment patterns. UL sites were strongly enriched for liver function regulators, including PPAR-α (p = 10^−83^), a key metabolic regulator, and C/EBPβ (p = 10^−54^), which regulates liver regeneration (**Supp. Table 5**). Analysis using published ChIP-seq data ^33–40^ validated these findings. The liver TFs FoxA1, C/EBPβ, and HNF4α, as well as the fasting modulator PPAR-α, all showed enrichment in UL, but not UH (**Fig. 3A-D, Supp. Fig. 1A-D**). These patterns parallel previous findings in human liver and hepatocytes ^5,21,41,42^. UH sites showed enrichment for KLF transcription factor family motifs (p = 10^−17^), proteins crucial for liver physiology and pathology ^43–47^ (**Supp. Table 6**).

**Figure 3:**
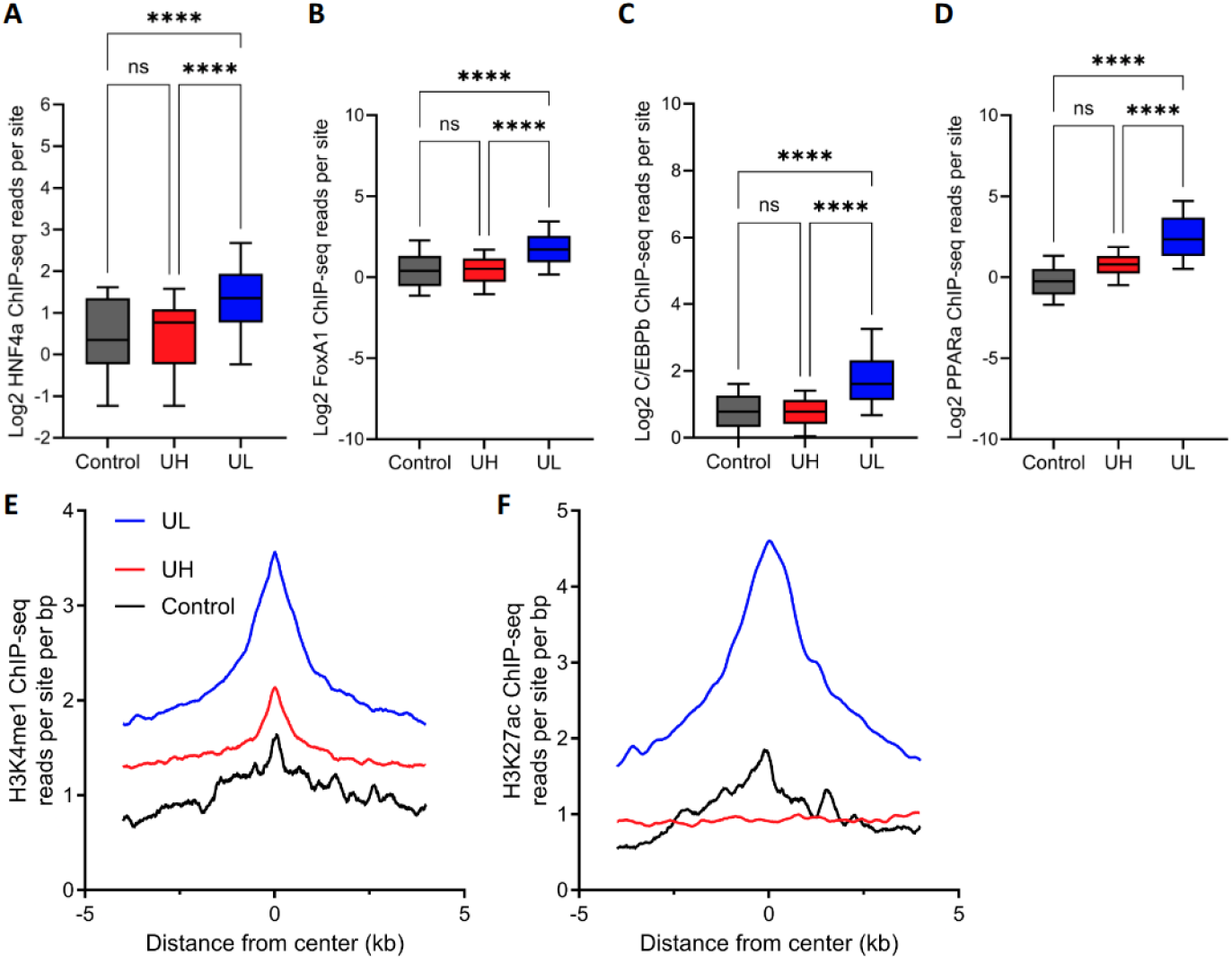
UL sites are bound by liver TFs and enhancer histone modifications. ChIP-seq analysis comparing the reads per site of UL, UH and controls for HNF4α (**A**), FoxA1 (**B**), C/EBPβ (**C**), and PPAR-α (**D**). **** p-value < 0.0001. H3K4me1 (**E**) and H3K27ac (**F**) ChIP-seq signal at a 5kb window relative to UL, UH and control site.

To enable a comprehensive chromatin-state analysis, we generated an additional list of mouse liver-unique methylation sites (**Supp. Table 7**), using the smaller mammalian methylation array ^10,48^. Chromatin-state analysis using eForge 2 ^49^ identified that like in humans ^5^, UL sites showed strong enrichment for liver-specific enhancers (**Supp. Table 8**). UH sites were enriched for both enhancers and areas flanking active TSS, but with significantly lower p-values and with no tissue specificity (**Supp. Table 9**). Indeed, ChIP-seq analysis of liver samples identified UL sites as enriched for the enhancer histone modifications H3K27ac and H3K4me1 (**Fig. 3E,F**). By contrast, UH sites showed no enrichment in H3K27ac, and a weak enrichment over background for H3K4me1 (**Fig. 3E,F**). Importantly, analysis of other tissues revealed the exact opposite. UH sites were enriched for both H3K27ac and H3K4me1, while UL sites showed no enrichment (**Supp. Fig. 1E-H**). Thus we concluded that UL sites significantly overlap tissue-specific enhancers.

**Supplementary Figure 1:**
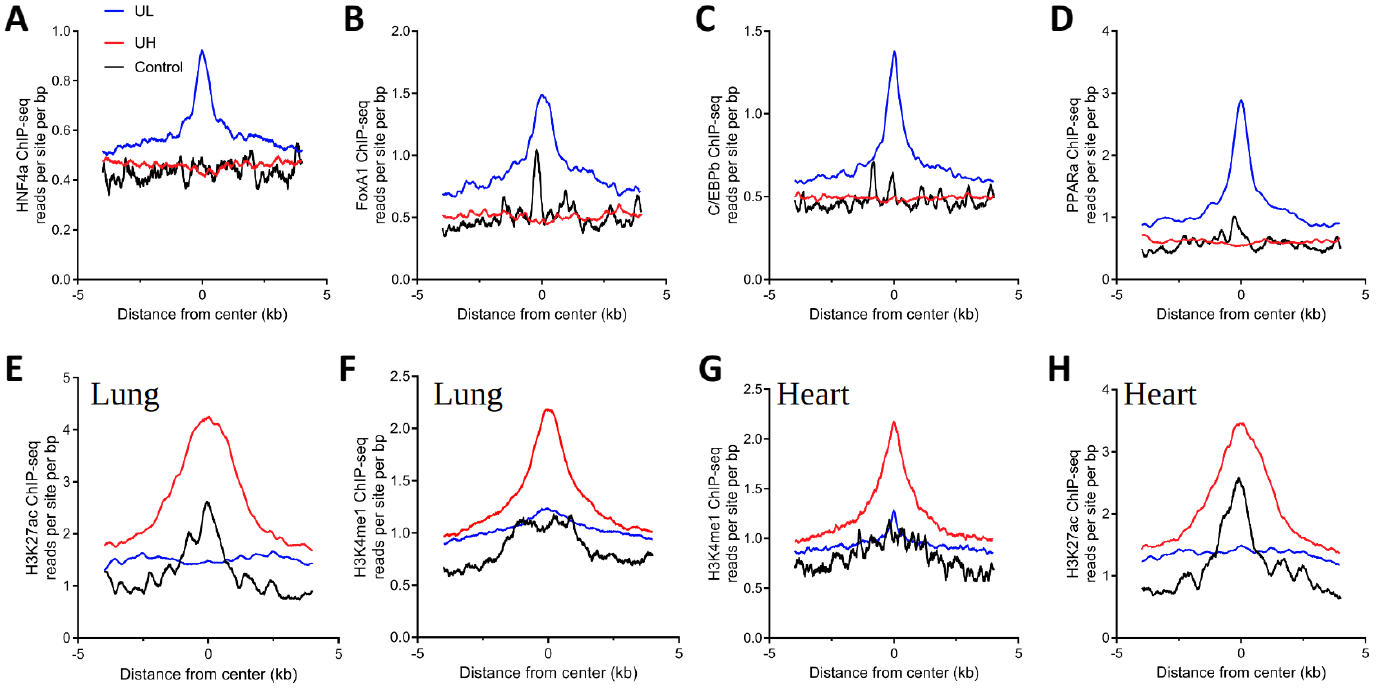
ChIP-seq read mapping relative to UL and UH sites. Liver ChIP-seq signal at a 5kb window relative to UL, UH and control site for HNF4α (**A**), FoxA1 (**B**), C/EBPβ (**C**), PPAR-α (**D**). H3K4me1 (**E**) and H3K27ac (**F**) ChIP-seq signal at a 5kb window relative to UL, UH and control site in lung tissue. H3K4me1 (**G**) and H3K27ac (**H**) ChIP-seq signal at a 5kb window relative to UL, UH and control site in heart tissue.

### Liver-unique methylation sites are enriched for aging-causal sites

To distinguish between causal and correlative relationships, we examined overlap with methylation clocks where causality was established through epigenome-wide Mendelian randomization analysis ^9^. We analyzed two such clocks: DamAge and AdaptAge, which track detrimental and adaptive methylation changes during aging, respectively. DamAge showed strong enrichment specifically for UH sites (p = 2 × 10^−5^), and all these sites carried positive weights in the clock (p = 0.01), indicating their contribution to accelerated aging. In contrast, AdaptAge showed only marginal enrichment (p = 0.046). UL sites showed no enrichment in either clock and were depleted in DamAge (0 sites found; p = 0.009; **Fig. 4A**). These findings suggest that the loss of epigenetic information at UH sites may causally contribute to liver aging.

**Figure 4:**
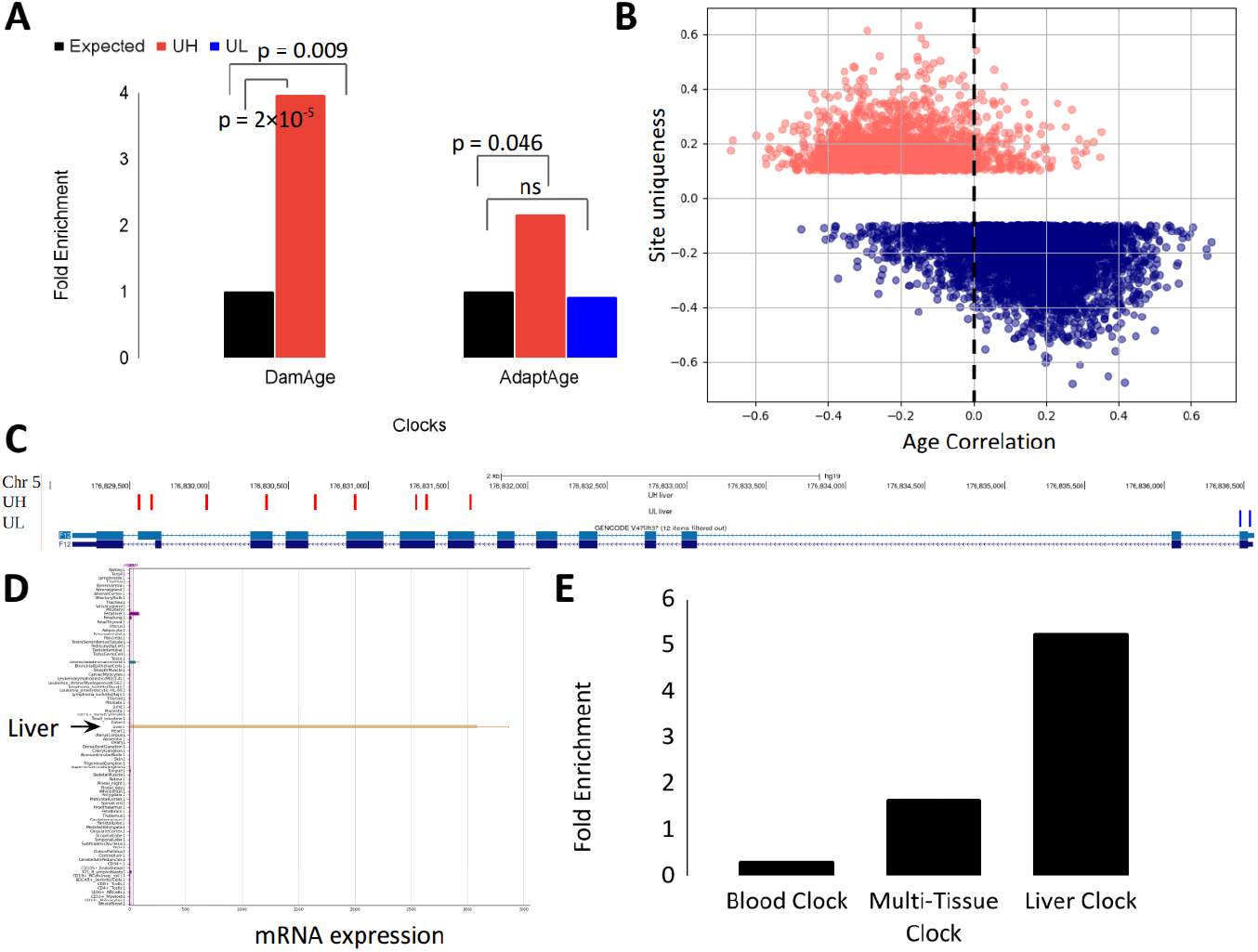
UH sites are enriched for causal-detrimental methylation sites. (**A**) Enrichment and depletion of UH and UL sites over expected values for the DamAge and AdaptAge clocks. (**B**) Methylation site age correlation plotted against site “uniqueness” (see Methods) demonstrated that the majority of UH sites showed a negative correlation, while UL sites showed a positive one, indicating regression to the mean upon aging. (**C**) Genome-browser view of liver-unique methylation sites along human coagulation factor XII, transcribed from right to left (**D**) Human F12 is expressed exclusively in the liver, as viewed by bioGPS, using the GeneAtlas U133A, gcrma dataset ^50,51^. The weaker magenta bar is the fetal liver. (**E**) Mouse liver-unique methylation sites are enriched in the mouse multi-tissue and liver clocks, but not in the blood clock.

### Liver-unique methylation sites regress to the mean upon aging

To test for potential overlap between age-associated changes in methylation and liver-unique methylation sites, we plotted the site “uniqueness” (see Methods) vs. correlation to aging. UH sites have an average negative correlation to aging, while UL sites have an average positive correlation (**Fig. 4B**). Thus, the majority of sites regress to the mean upon aging. Interestingly, no obvious relation between the site “uniqueness” and age-correlation was observed. This is in contrast to a more quantitative relation observed for example in kidney-unique sites in CKD ^18^.

### Genes associated with liver-unique methylation sites in mice and human show limited overlap

We note that UL and UH should not simply be interpreted as expressed and repressed genes, respectively. The role of methylation in gene regulation is context dependent ^52,53^. For example, both low methylation levels adjacent to the TSS and high methylation levels along the gene body positively correlate with expression level. This pattern is observed in the human coagulation factor XII (F12), predominantly expressed in the liver, where UL sites are found in the TSS, while UH sites are scattered across the gene body and later exons (**Fig. 4C, D**).

We compared genes associated with liver-specific methylation sites in mice (N = 2177) and humans (N = 1611), observing a 17% match in mice and a 23% match in humans. While statistically significant (p = 10^−5^), this overlap appears modest. We thus tested if a significant overlap exists on the pathway level using the STRING database ^54^. This analysis identified several common pathways enriched in both species. Notably, both human and mouse analyses showed enrichment in Metabolic pathways (human: FDR < 10^−6^, mouse: FDR < 10^−7^) and Complement and coagulation (human: FDR < 10^−10^, mouse: FDR = 0.00026). These results can reflect on processes conserved across species particularly in the context of liver sole function in several metabolic processes and coagulation cascade regulation.

### Liver-unique methylation sites are enriched in epigenetic clocks in mice

To explore the relationship between epigenetic information loss and aging, we first examined mouse DNA methylation clocks ^12^, which accurately measure chronological age and may capture aspects of biological aging. Comparison of liver-unique methylation sites with the mouse blood clock demonstrated no enrichment (p = 0.25; **Fig. 4E)**. A multi-tissue clock revealed a modest and statistically non-significant 1.7-fold enrichment (p = 0.1; **Fig. 4E)**. However, a mouse liver-specific clock showed a 5.3-fold enrichment (p = 0.0001; **Fig. 4E**). These results suggest that liver-unique methylation sites are relevant to aging in a tissue-specific manner (**Supp. Table 10**). Interestingly, both UH and UL sites were represented in these clocks at ratios proportional to their overall prevalence in the liver methylome.

### Repeated fasting partially restores epigenetic information loss in mice livers

DNA methylation patterns can change due to environmental effects. Moreover, both beneficial behaviours and recovery from disease can result in temporal “reversion” of at least some epigenetic clocks ^12,55–58^. However, while the reversibility of epigenetic information loss was established with exogenous interventions like treatment with “Yamanaka Factors”, its reversibility under natural settings has been underexplored. We note that this is distinguished from exploring the reversibility of various epigenetic clocks that have been studied more carefully ^56^. To investigate the potential reversibility of epigenetic information loss, we examined liver DNA accessibility in mice subjected to alternate-day fasting (ADF). ADF, a form of intermittent fasting, reduces liver pathologies and enhances enhancer sensitivity ^59–61^. We analyzed DNA accessibility in alternate-day fasting mice vs. mice subjected to unrestricted feeding (^59^; **Supp. Table 11**). Enhancers showing increased accessibility were 8-fold enriched in liver-unique sites vs. controls (**Fig. 5A**; **Supp. Table 11**). This effect was exclusively driven by UL sites that showed a 12-fold enrichment vs. controls and a 44-fold enrichment vs. UH (**Fig. 5B**; **Supp. Table 11**). Enhancers showing decreased accessibility were 4-fold enriched in liver-unique sites vs. controls (**Fig. 5A**), however, this effect was driven by UH sites that were 7-fold enriched vs. controls and 3-fold enriched vs. UL (**Fig. 5C**). This phenomenon was not observed when comparing livers of mice subjected to a single 24-hour fast (**Fig. 5A-C**). These findings demonstrate that liver epigenetic information lost can be partially restored through repeated fasting (**Fig. 5D**), suggesting a potential therapeutic approach for reversing tissue-specific epigenetic deterioration.

**Figure 5:**
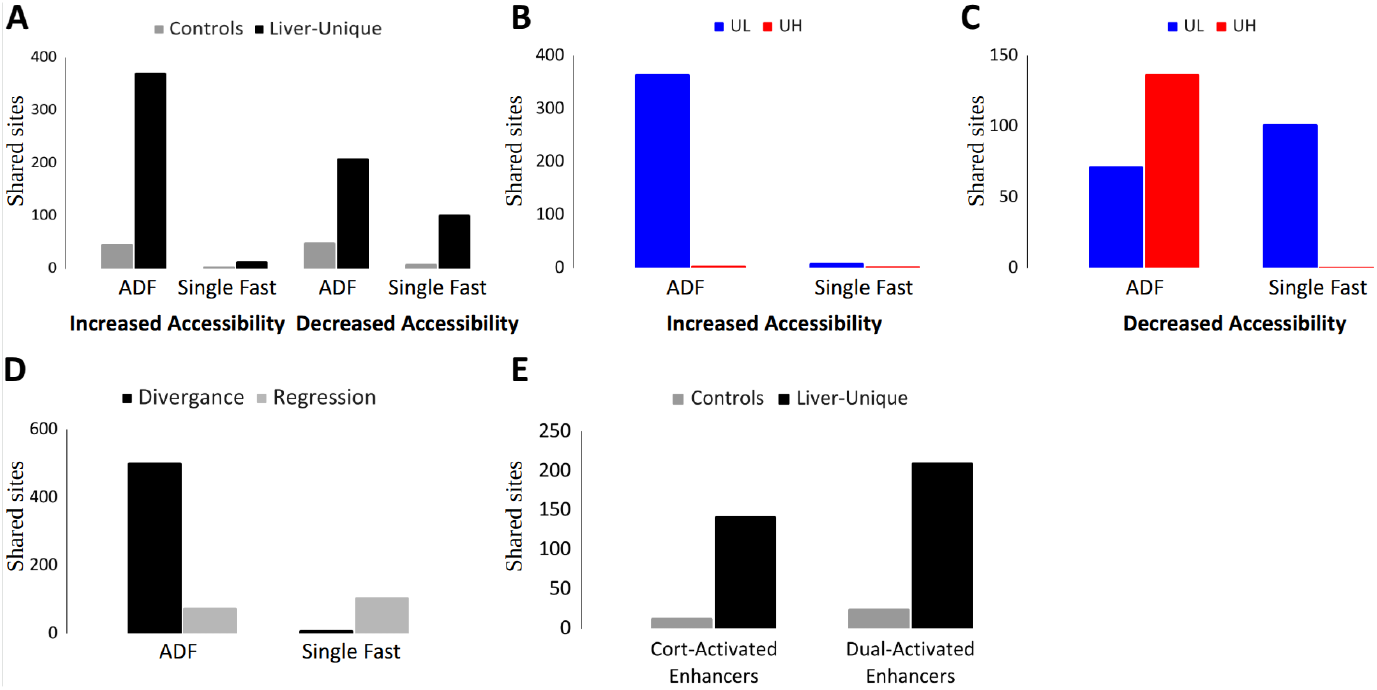
Reversibility of epigenetic information loss in the liver of fasting mice. (**A**) Overlap of liver-unique methylation sites and random controls with enhancers that show increased (left) or decreased (right) accessibility following alternate-day fasting (ADF) or a single fast, shows a greater overlap upon ADF. (**B**) The overlap in increased accessibility enhancers is driven exclusively by UL sites in ADF. (**C**) The overlap in decreased accessibility enhancers is mostly driven by UH sites in ADF, but some UL sites change following ADF and a single fast. (**D**) Divergence from the mean and regression to the mean in liver unique sites following ADF or a single fast. (**E**) Overlap of liver-unique methylation sites with corticosterone-activated and dual-activated enhancers.

### Epigenetic information is restored in hepatocytes upon fasting mimicry

While bulk-tissue analysis reveals epigenetic information loss and restoration, it does not inform us if these changes are driven by changes in cell composition, changes within cells or both. To address this limitation, we leveraged an *ex vivo* system employed by Goldberg et al. ^62^ that mimics long-term fasting in isolated hepatocytes through agonist treatment - corticosterone to activate the glucocorticoid receptor and Wy-14643 to activate PPARα. Analysis of liver-unique methylation sites revealed a striking 10-fold enrichment in corticosterone-activated enhancers compared to controls (143 enhancers versus 14 in random controls; **Fig. 5E**; **Supp. Table 12**). Remarkably, all identified enhancers (143/143) mapped to UL sites (p < 10^−29^). These findings were further validated using dual enhancers identified through combined activation with Wy-14643 and corticosterone, where an 8-fold enrichment was observed (211 vs. 26; **Fig. 5E**; **Supp. Table 12**). Again, nearly all identified enhancers (210/211; p < 10^−40^) mapped to UL sites. Together, these results demonstrate that epigenetic information loss and restoration occurs at the cellular level within hepatocytes.

## Discussion

Our findings establish the reversibility of epigenetic information loss in liver aging and disease, while demonstrating its causal role in these processes. The regression of liver-unique methylation patterns toward pan-tissue averages during cirrhosis validates and extends previous observations of epigenetic information loss in liver pathologies. Importantly, the strong enrichment of liver-unique methylation sites near disease-associated genetic variants suggests a mechanistic link between genetic predisposition to liver disease and tissue-specific epigenetic regulation.

The distinct behavior of UH and UL sites in aging presents a nuanced picture of epigenetic information loss. Both UH and UL sites show age-correlated methylation changes and epigenetic information loss, revealed through correlation analysis of methylation patterns in mouse liver samples. This association was further strengthened by Mendelian randomization analysis showing significant enrichment of human UH sites in the DamAge clock, with all identified sites carrying positive weights, indicating their contribution to accelerated aging. The causal nature of these changes is particularly notable given the modest enrichment of UH in the AdaptAge clock, which tracks adaptive age-related changes. Conversely, UL sites are depleted in DamAge and showed no enrichment in AdaptAge, despite their clear susceptibility to disease-associated changes, as demonstrated in our cirrhosis analysis. Together, these findings suggest a model where UH sites actively participate in the aging process, possibly due to their importance in other tissues, while UL sites primarily respond to disease states.

The conservation of pathway-level associations between mouse and human liver-unique methylation sites, despite limited gene-level overlap, points to the fundamental role of these epigenetic patterns in liver function. The enrichment of metabolic and coagulation pathways in both species aligns with core liver functions, while species-specific enrichments may reflect a lower level of conservation, evolutionary adaptations, or experimental constraints.

Perhaps most significantly, our demonstration that repeated fasting can partially restore liver-unique methylation patterns provides evidence for the reversibility of epigenetic information loss through physiological interventions. The cell-autonomous nature of this restoration, as shown in isolated hepatocytes under fasting-mimicking conditions, indicates that these changes occur at the cellular level rather than through alterations in tissue composition. This finding has important implications for understanding the nature and reversibility of disease and age related epigenetic changes.

Several limitations of this study should be considered. While our findings demonstrate the reversibility of epigenetic information loss in mouse models, the extent to which these results translate to humans remains to be determined. Our analysis of human samples was limited to cross-sectional data, preventing direct assessment of temporal dynamics and individual variation in epigenetic changes. The cell-autonomous effects observed in isolated hepatocytes, while promising, were studied under artificial fasting-mimicking conditions that may not fully recapitulate the complexity of *in vivo* fasting responses. Additionally, while we show enrichment of liver-unique methylation sites near disease-associated variants, the precise molecular mechanisms linking these epigenetic patterns to disease progression remain unclear. One significant limitation is our intermittent use of DNA methylation and accessibility data. Although differential DNA accessibility provides both a functional readout and a strong indicator of methylation profile changes, direct measurement of DNA methylation levels in the fasting intervention studies would strengthen our conclusions. Finally, while we demonstrate partial restoration of epigenetic information through fasting, the long-term stability of these changes and their impact on liver function and disease outcomes requires further investigation in longitudinal studies.

These results advance our understanding of epigenetic information loss as both a marker and mediator of aging and disease, while offering promising directions for therapeutic interventions. Future studies should investigate the molecular mechanisms linking metabolic interventions to epigenetic restoration and explore the potential of targeted approaches to reverse tissue-specific epigenetic deterioration in aging and disease.

## Materials and methods

### Identification of mouse liver-unique sites

Liver-unique methylation sites in mice were generated using methods similar to ^5^, based on public data from GSE184410 ^31,32^. 92 *Mus musculus* liver samples were selected, along with 404 *Mus musculus* samples from 19 other tissues as the background. A unique high methylation site in the liver is defined as a site where the average methylation value is higher by 0.1 or more than the 0.95 quantile of all other tissues, and a unique low methylation site is one where the average methylation value is lower by 0.1 or more than the 0.05 quantile of all other tissues. Similarly, we’ve extracted mouse liver unique sites from 4,011 Mus musculus liver samples found in GSE223748 ^10^, generated by the Mammalian Methylation Consortium using the Mammal40k array ^48^.

### CpG sites correlation with age

Pearson correlation between age and methylation levels was plotted for all mouse liver-unique CpG sites. Data was sourced from the GSE184410 database, including all liver samples with no pathologies.

### Statistics

Unless otherwise noted, p-values were calculated using log-scale binomial or hypergeometric distribution on R. In the cirrhosis overlaps (Fig. 1), pathogenic SNP analysis (Fig. 2), and mouse epigenetic clocks (Fig. 4), the p-values were determined using a Monte Carlo simulation with 10,000 iterations. Control groups of the same size as the liver-unique sites (N = 3,674 for human; N = 6,039 for mouse) were randomly sampled from tissue-unique sites excluding liver-unique ones (N = 84,249 for human; N = 25,128 for mouse ^5^). When the observed intersection of liver-unique sites exceeded all values from the simulated distribution, the p-value was calculated as 1/number of simulations, yielding a p-value of 0.0001.

### Enrichment analysis

Enrichment analysis for cirrhosis, DamAge, AdaptAge, ADF and hepatocytes was done using randomly selected controls as described above, requiring either an identical CpG site or an overlap between the sites and range. For pathogenic SNPs and mouse tissue clocks, sites considered overlapping if found below a distance threshold, found in Supp. Table 3 for the SNPs, or below 2.5kb for the clocks. Reducing this threshold did not change the enrichment of the liver clock, but reduced the number of overlapping sites.

### Epigenetic state analysis

Chromatin state analysis was performed on Liver-unique methylation sites in mice using the eFORGE ^63^ web server (https://eforge40k.altiusinstitute.org/). It was set to eFORGE 40K (mammalian array version), “Consolidated Roadmap Epigenomics—Chromatin—All 15 state marks” and default setting of 1 kb window, 1000 background repetition, 0.01 strict and 0.05 marginal.

### Motif enrichment analysis

Known and de novo motifs were identified and enrichment was calculated using the Hypergeometric Optimization of Motif EnRichment (HOMER; ^64^). Analysis was done separately for mouse UL and UH sites, analyzing sequences extracted from the mm10 genome assembly and spanning ±500bp around the methylation probe start sites^65^. The Infinium mouse methylation manifest file served as the background for motif enrichment analysis.

### ChIP-seq analysis

Tag density of ChIP signal around methylated sites (±100 bp) were analyzed using the HOMER suite. In aggregate plots, the tag count (averaged across all sites) per site per base pair was calculated using the HOMER suite (annotatePeaks, option -size 8000 -hist 10). In box plots, tag count ±200 bp around the site (averaged across all sites) was calculated using the HOMER suite (annotatePeaks, option -size 400 -noann). In box plots for H3K27ac, which is a broader signal, tag count ±500 bp around the site (averaged across all sites) was calculated using the HOMER suite (annotatePeaks, option -size 1000 -noann). In both aggregate plots and box plots, the data is an average of all available replicates. In all box plots, the 10–90 percentiles are plotted.

### Data and materials availability

Generated data are available as supplementary files.

## Acknowledgments

We thank the Bar lab members for comments and suggestions. This work was supported by the Israeli Science Foundation (grants 654/20 and 632/20 to DZB), the Center for Artificial Intelligence & Data Science in Tel Aviv University (TAD to DZB) and the European Research Council (grant 947907 to IG).

## Author contributions

DZB, NS and RBS designed the experiments. RBS and NS performed the analysis. NK performed the ChIP-seq analysis. DZB wrote the manuscript with critical inputs and comments from NS, RBS, NK, IG and MG.

