## Supplementary material for "Reversible and Causal Epigenetic Information Loss in Liver Aging and Disease": Supp. Table 5

| Rank | Motif | Name | P-value | log P-pvalue | q-value (Benjamini) | # Target Sequences with Motif | % of Targets Sequences with Motif | # Background Sequences with Motif | % of Background Sequences with Motif | Motif File | SVG |
|---|---|---|---|---|---|---|---|---|---|---|---|
| 1 | 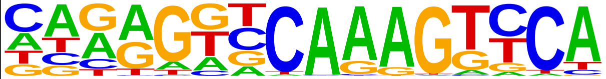 | HNF4a(NR),DR1/HepG2-HNF4a-ChIP-Seq(GSE25021)/Homer | 1e-66 | -1.521e+02 | 0.0000 | 1335.0 | 35.41% | 65605.6 | 22.98% | motif file (matrix) | svg |
| 2 | 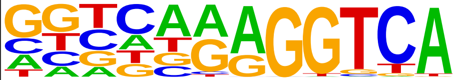 | COUP-TFII(NR)/K562-NR2F1-ChIP-Seq(Encode)/Homer | 1e-55 | -1.276e+02 | 0.0000 | 2570.0 | 68.17% | 158731.0 | 55.61% | motif file (matrix) | svg |
| 3 | 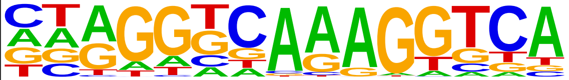 | PPARa(NR),DR1/Liver-Ppara-ChIP-Seq(GSE47954)/Homer | 1e-52 | -1.215e+02 | 0.0000 | 2157.0 | 57.21% | 127666.0 | 44.73% | motif file (matrix) | svg |
| 4 | 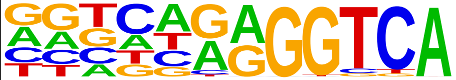 | EAR2(NR)/K562-NR2F6-ChIP-Seq(Encode)/Homer | 1e-50 | -1.154e+02 | 0.0000 | 2414.0 | 64.03% | 148292.8 | 51.95% | motif file (matrix) | svg |
| 5 | 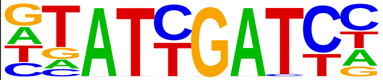 | HNF6(Homeobox)/Liver-Hnf6-ChIP-Seq(ERP000394)/Homer | 1e-48 | -1.118e+02 | 0.0000 | 1243.0 | 32.97% | 64194.2 | 22.49% | motif file (matrix) | svg |
| 6 | 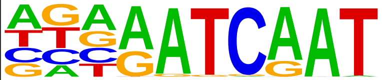 | Cux2(Homeobox)/Liver-Cux2-ChIP-Seq(GSE35985)/Homer | 1e-47 | -1.098e+02 | 0.0000 | 1061.0 | 28.14% | 52520.0 | 18.40% | motif file (matrix) | svg |
| 7 | 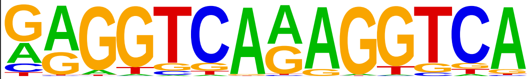 | TR4(NR),DR1/Hela-TR4-ChIP-Seq(GSE24685)/Homer | 1e-43 | -9.942e+01 | 0.0000 | 438.0 | 11.62% | 16257.0 | 5.70% | motif file (matrix) | svg |
| 8 | 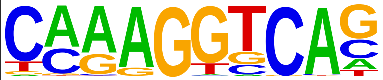 | Erra(NR)/HepG2-Erra-ChIP-Seq(GSE31477)/Homer | 1e-42 | -9.897e+01 | 0.0000 | 3124.0 | 82.86% | 209366.8 | 73.35% | motif file (matrix) | svg |
| 9 | 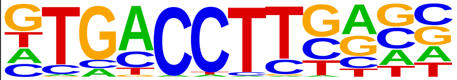 | ERRg(NR)/Kidney-ESRRG-ChIP-Seq(GSE104905)/Homer | 1e-41 | -9.576e+01 | 0.0000 | 1774.0 | 47.06% | 103421.0 | 36.23% | motif file (matrix) | svg |
| 10 | 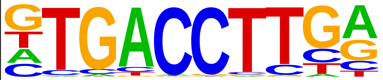 | Esrrb(NR)/mES-Esrrb-ChIP-Seq(GSE11431)/Homer | 1e-41 | -9.565e+01 | 0.0000 | 1538.0 | 40.80% | 86618.1 | 30.35% | motif file (matrix) | svg |
| 11 | 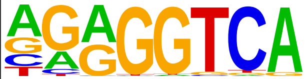 | COUP-TFII(NR)/Artia-Nr2f2-ChIP-Seq(GSE46497)/Homer | 1e-37 | -8.525e+01 | 0.0000 | 2701.0 | 71.64% | 176157.2 | 61.71% | motif file (matrix) | svg |
| 12 | 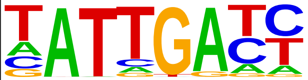 | Hnf6b(Homeobox)/LNCaP-Hnf6b-ChIP-Seq(GSE106305)/Homer | 1e-35 | -8.134e+01 | 0.0000 | 1694.0 | 44.93% | 100052.9 | 35.05% | motif file (matrix) | svg |
| 13 | 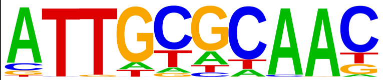 | CEBP(bZIP)/ThioMac-CEBPb-ChIP-Seq(GSE21512)/Homer | 1e-33 | -7.665e+01 | 0.0000 | 1307.0 | 34.67% | 73520.3 | 25.76% | motif file (matrix) | svg |
| 14 | 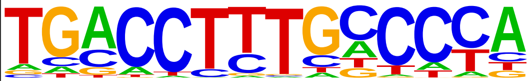 | PPARE(NR),DR1/3T3L1-Pparg-ChIP-Seq(GSE13511)/Homer | 1e-32 | -7.577e+01 | 0.0000 | 1859.0 | 49.31% | 113074.8 | 39.61% | motif file (matrix) | svg |

| | | | | | | | | | | | |
|---|---|---|---|---|---|---|---|---|---|---|---|
| 15 | 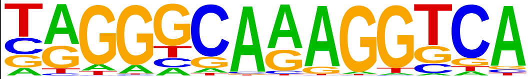 | RXR(NR),DR1/3T3L1-RXR-ChIP-Seq(GSE13511)/Homer | 1e-30 | -7.029e+01 | 0.0000 | 2044.0 | 54.22% | 127881.9 | 44.80% | motif file (matrix) | svg |
| 16 | 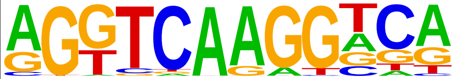 | RAR:RXR(NR),DR0/ES-RAR-ChIP-Seq(GSE56893)/Homer | 1e-27 | -6.327e+01 | 0.0000 | 546.0 | 14.48% | 25607.1 | 8.97% | motif file (matrix) | svg |
| 17 | 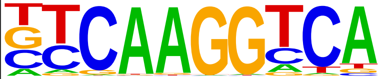 | Nr5a2(NR)/Pancreas-LRH1-ChIP-Seq(GSE34295)/Homer | 1e-27 | -6.299e+01 | 0.0000 | 1613.0 | 42.79% | 97593.4 | 34.19% | motif file (matrix) | svg |
| 18 | 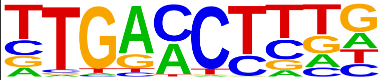 | RARa(NR)/K562-RARa-ChIP-Seq(Encode)/Homer | 1e-26 | -6.138e+01 | 0.0000 | 3436.0 | 91.14% | 243482.9 | 85.30% | motif file (matrix) | svg |
| 19 | 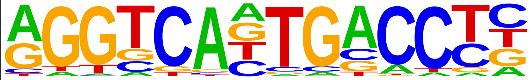 | FXR(NR),IR1/Liver-FXR-ChIP-Seq(Chong_et_al.)/Homer | 1e-25 | -5.913e+01 | 0.0000 | 868.0 | 23.02% | 46642.1 | 16.34% | motif file (matrix) | svg |
| 20 | 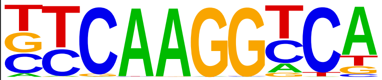 | Nr5a2(NR)/mES-Nr5a2-ChIP-Seq(GSE19019)/Homer | 1e-24 | -5.608e+01 | 0.0000 | 1286.0 | 34.11% | 75676.9 | 26.51% | motif file (matrix) | svg |
| 21 | 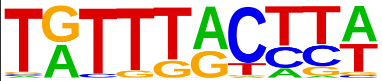 | FOXM1(Forkhead)/MCF7-FOXM1-ChIP-Seq(GSE72977)/Homer | 1e-24 | -5.548e+01 | 0.0000 | 2108.0 | 55.92% | 135829.9 | 47.59% | motif file (matrix) | svg |
| 22 | 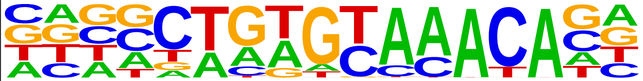 | Fox:Ebox(Forkhead,bHLH)/Panc1-Foxa2-ChIP-Seq(GSE47459)/Homer | 1e-22 | -5.087e+01 | 0.0000 | 2066.0 | 54.80% | 133710.6 | 46.84% | motif file (matrix) | svg |
| 23 | 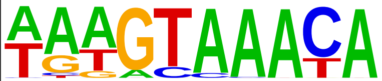 | FOXA1(Forkhead)/MCF7-FOXA1-ChIP-Seq(GSE26831)/Homer | 1e-20 | -4.657e+01 | 0.0000 | 2019.0 | 53.55% | 131194.1 | 45.96% | motif file (matrix) | svg |
| 24 | 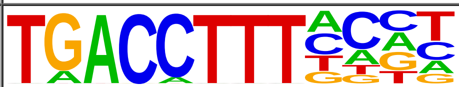 | Nur77(NR)/K562-NR4A1-ChIP-Seq(GSE31363)/Homer | 1e-18 | -4.322e+01 | 0.0000 | 500.0 | 13.26% | 25226.8 | 8.84% | motif file (matrix) | svg |
| 25 | 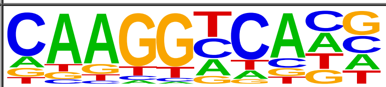 | SF1(NR)/H295R-Nr5a1-ChIP-Seq(GSE44220)/Homer | 1e-18 | -4.317e+01 | 0.0000 | 1144.0 | 30.34% | 68341.3 | 23.94% | motif file (matrix) | svg |
| 26 | 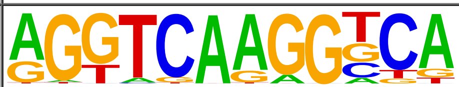 | RARg(NR)/ES-RARg-ChIP-Seq(GSE30538)/Homer | 1e-18 | -4.297e+01 | 0.0000 | 296.0 | 7.85% | 12904.2 | 4.52% | motif file (matrix) | svg |
| 27 | 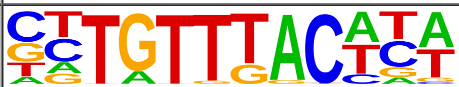 | Foxa2(Forkhead)/Liver-Foxa2-ChIP-Seq(GSE25694)/Homer | 1e-18 | -4.167e+01 | 0.0000 | 1660.0 | 44.03% | 105717.8 | 37.04% | motif file (matrix) | svg |
| 28 | 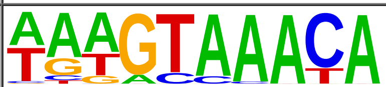 | FOXA1(Forkhead)/LNCAP-FOXA1-ChIP-Seq(GSE27824)/Homer | 1e-17 | -3.963e+01 | 0.0000 | 2254.0 | 59.79% | 150863.1 | 52.85% | motif file (matrix) | svg |
| 29 | 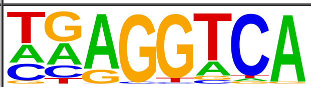 | THRb(NR)/Liver-NR1A2-ChIP-Seq(GSE52613)/Homer | 1e-14 | -3.450e+01 | 0.0000 | 3647.0 | 96.74% | 267982.3 | 93.88% | motif file (matrix) | svg |
| 30 | 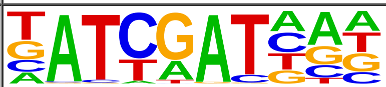 | CUX1(Homeobox)/K562-CUX1-ChIP-Seq(GSE92882)/Homer | 1e-14 | -3.406e+01 | 0.0000 | 1279.0 | 33.93% | 80027.3 | 28.04% | motif file (matrix) | svg |

| | | | | | | | | | | | |
|---|---|---|---|---|---|---|---|---|---|---|---|
| 31 | 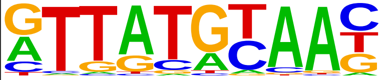 | HLF(bZIP)/HSC-HLF.Flag-ChIP-Seq(GSE69817)/Homer | 1e-14 | -3.272e+01 | 0.0000 | 1494.0 | 39.63% | 95923.5 | 33.61% | motif file (matrix) | svg |
| 32 | 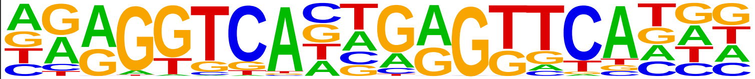 | VDR(NR),DR3/GM10855-VDR+vitD-ChIP-Seq(GSE22484)/Homer | 1e-12 | -2.914e+01 | 0.0000 | 483.0 | 12.81% | 26292.4 | 9.21% | motif file (matrix) | svg |
| 33 | 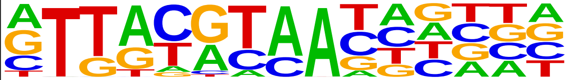 | NFIL3(bZIP)/HepG2-NFIL3-ChIP-Seq(Encode)/Homer | 1e-12 | -2.877e+01 | 0.0000 | 1169.0 | 31.01% | 73559.6 | 25.77% | motif file (matrix) | svg |
| 34 | 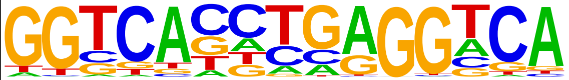 | THRb(NR)/HepG2-THRb.Flag-ChIP-Seq(Encode)/Homer | 1e-11 | -2.586e+01 | 0.0000 | 1061.0 | 28.14% | 66666.3 | 23.36% | motif file (matrix) | svg |
| 35 | 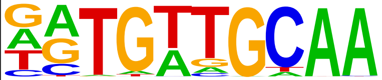 | CEBP:AP1(bZIP)/ThioMac-CEBPb-ChIP-Seq(GSE21512)/Homer | 1e-10 | -2.330e+01 | 0.0000 | 1338.0 | 35.49% | 87367.6 | 30.61% | motif file (matrix) | svg |
| 36 | 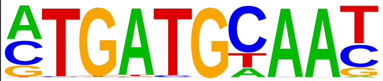 | Atf4(bZIP)/MEF-Atf4-ChIP-Seq(GSE35681)/Homer | 1e-9 | -2.207e+01 | 0.0000 | 543.0 | 14.40% | 31651.3 | 11.09% | motif file (matrix) | svg |
| 37 | 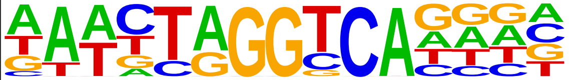 | RORa(NR)/Liver-Rora-ChIP-Seq(GSE101115)/Homer | 1e-9 | -2.123e+01 | 0.0000 | 341.0 | 9.05% | 18453.8 | 6.46% | motif file (matrix) | svg |
| 38 | 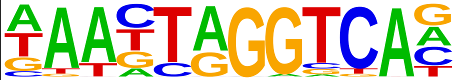 | RORg(NR)/Liver-Rorc-ChIP-Seq(GSE101115)/Homer | 1e-8 | -2.001e+01 | 0.0000 | 231.0 | 6.13% | 11667.1 | 4.09% | motif file (matrix) | svg |
| 39 | 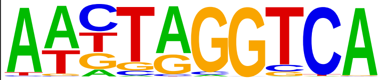 | RORgt(NR)/EL4-RORgt.Flag-ChIP-Seq(GSE56019)/Homer | 1e-8 | -1.977e+01 | 0.0000 | 310.0 | 8.22% | 16706.5 | 5.85% | motif file (matrix) | svg |
| 40 | 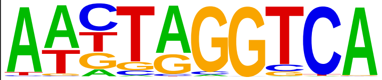 | RORgt(NR)/EL4-RORgt.Flag-ChIP-Seq(GSE56019)/Homer | 1e-8 | -1.977e+01 | 0.0000 | 310.0 | 8.22% | 16706.5 | 5.85% | motif file (matrix) | svg |
| 41 | 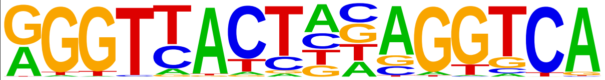 | LXRE(NR),DR4/RAW-LXRb.biotin-ChIP-Seq(GSE21512)/Homer | 1e-8 | -1.943e+01 | 0.0000 | 138.0 | 3.66% | 6139.9 | 2.15% | motif file (matrix) | svg |
| 42 | 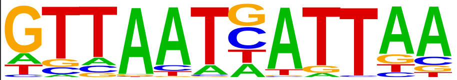 | HNF1b(Homeobox)/PDAC-HNF1B-ChIP-Seq(GSE64557)/Homer | 1e-8 | -1.900e+01 | 0.0000 | 365.0 | 9.68% | 20425.5 | 7.16% | motif file (matrix) | svg |
| 43 | 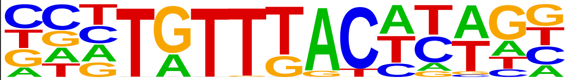 | Foxa3(Forkhead)/Liver-Foxa3-ChIP-Seq(GSE77670)/Homer | 1e-8 | -1.900e+01 | 0.0000 | 758.0 | 20.11% | 47230.2 | 16.55% | motif file (matrix) | svg |
| 44 | 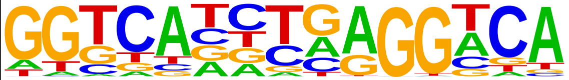 | THRa(NR)/C17.2-THRa-ChIP-Seq(GSE38347)/Homer | 1e-7 | -1.786e+01 | 0.0000 | 806.0 | 21.38% | 50944.8 | 17.85% | motif file (matrix) | svg |
| 45 | 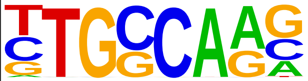 | NF1-halfsite(CTF)/LNCaP-NF1-ChIP-Seq(Unpublished)/Homer | 1e-7 | -1.748e+01 | 0.0000 | 2504.0 | 66.42% | 177362.2 | 62.14% | motif file (matrix) | svg |
| 46 | 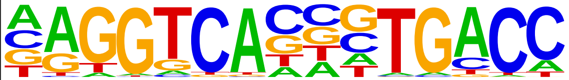 | ERE(NR),IR3/MCF7-ERa-ChIP-Seq(Unpublished)/Homer | 1e-7 | -1.730e+01 | 0.0000 | 591.0 | 15.68% | 36086.2 | 12.64% | motif file (matrix) | svg |

| | | | | | | | | | | | |
|---|---|---|---|---|---|---|---|---|---|---|---|
| 47 | 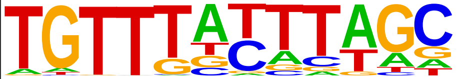 | FoxD3(forkhead)/ZebrafishEmbryo-Foxd3.biotin-ChIP-seq(GSE106676)/Homer | 1e-7 | -1.678e+01 | 0.0000 | 1707.0 | 45.28% | 116985.5 | 40.98% | motif file (matrix) | svg |
| 48 | 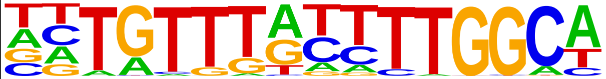 | NF1:FOXA1(CTF,Forkhead)/LNCAP-FOXA1-ChIP-Seq(GSE27824)/Homer | 1e-7 | -1.643e+01 | 0.0000 | 157.0 | 4.16% | 7610.9 | 2.67% | motif file (matrix) | svg |
| 49 | 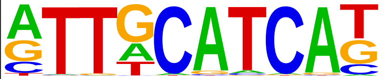 | Chop(bZIP)/MEF-Chop-ChIP-Seq(GSE35681)/Homer | 1e-6 | -1.547e+01 | 0.0000 | 418.0 | 11.09% | 24723.5 | 8.66% | motif file (matrix) | svg |
| 50 | 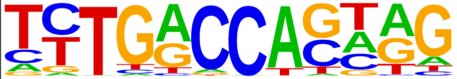 | Bcl11a(Zf)/HSPC-BCL11A-ChIP-Seq(GSE104676)/Homer | 1e-6 | -1.486e+01 | 0.0000 | 1295.0 | 34.35% | 87276.7 | 30.58% | motif file (matrix) | svg |
| 51 | 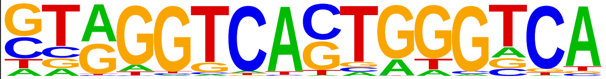 | Reverb(NR),DR2/RAW-Reverba.biotin-ChIP-Seq(GSE45914)/Homer | 1e-6 | -1.483e+01 | 0.0000 | 307.0 | 8.14% | 17439.5 | 6.11% | motif file (matrix) | svg |
| 52 | 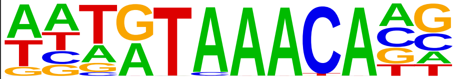 | FoxL2(Forkhead)/Ovary-FoxL2-ChIP-Seq(GSE60858)/Homer | 1e-6 | -1.388e+01 | 0.0000 | 1622.0 | 43.02% | 111899.7 | 39.20% | motif file (matrix) | svg |
| 53 | 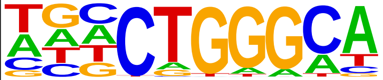 | ZNF416(Zf)/HEK293-ZNF416.GFP-ChIP-Seq(GSE58341)/Homer | 1e-5 | -1.376e+01 | 0.0000 | 2113.0 | 56.05% | 148952.1 | 52.18% | motif file (matrix) | svg |
| 54 | 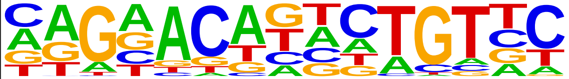 | PR(NR)/T47D-PR-ChIP-Seq(GSE31130)/Homer | 1e-5 | -1.232e+01 | 0.0000 | 3025.0 | 80.24% | 220473.0 | 77.24% | motif file (matrix) | svg |
| 55 | 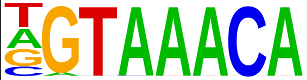 | Foxo3(Forkhead)/U2OS-Foxo3-ChIP-Seq(E-MTAB-2701)/Homer | 1e-5 | -1.155e+01 | 0.0001 | 1408.0 | 37.35% | 97092.7 | 34.01% | motif file (matrix) | svg |
| 56 | 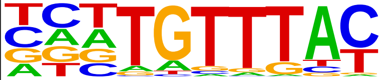 | FOXK1(Forkhead)/HEK293-FOXK1-ChIP-Seq(GSE51673)/Homer | 1e-4 | -1.144e+01 | 0.0001 | 1818.0 | 48.22% | 127771.7 | 44.76% | motif file (matrix) | svg |
| 57 | 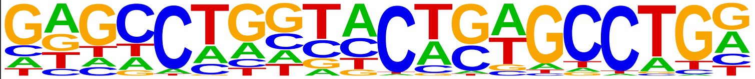 | ZNF322(Zf)/HEK293-ZNF322.GFP-ChIP-Seq(GSE58341)/Homer | 1e-4 | -1.128e+01 | 0.0001 | 567.0 | 15.04% | 36221.6 | 12.69% | motif file (matrix) | svg |
| 58 | 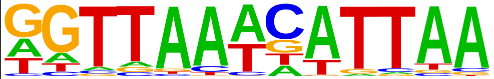 | Hnf1(Homeobox)/Liver-Foxa2-Chip-Seq(GSE25694)/Homer | 1e-4 | -1.045e+01 | 0.0002 | 312.0 | 8.28% | 18787.8 | 6.58% | motif file (matrix) | svg |
| 59 | 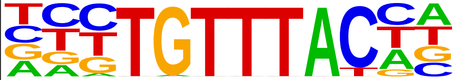 | FOXP1(Forkhead)/H9-FOXP1-ChIP-Seq(GSE31006)/Homer | 1e-4 | -9.997e+00 | 0.0003 | 902.0 | 23.93% | 60711.4 | 21.27% | motif file (matrix) | svg |
| 60 | 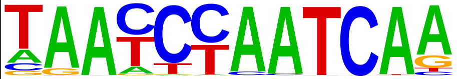 | Duxbl(Homeobox)/NIH3T3-Duxbl.HA-ChIP-Seq(GSE119782)/Homer | 1e-4 | -9.931e+00 | 0.0004 | 161.0 | 4.27% | 8850.6 | 3.10% | motif file (matrix) | svg |
| 61 | 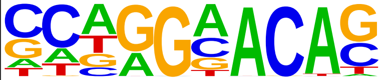 | AR-halfsite(NR)/LNCaP-AR-ChIP-Seq(GSE27824)/Homer | 1e-4 | -9.848e+00 | 0.0004 | 3588.0 | 95.17% | 267397.7 | 93.68% | motif file (matrix) | svg |
| 62 | 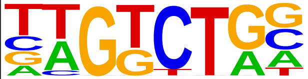 | Smad3(MAD)/NPC-Smad3-ChIP-Seq(GSE36673)/Homer | 1e-4 | -9.783e+00 | 0.0004 | 3303.0 | 87.61% | 243839.8 | 85.43% | motif file (matrix) | svg |

| | | | | | | | | | | | |
|---|---|---|---|---|---|---|---|---|---|---|---|
| 63 | 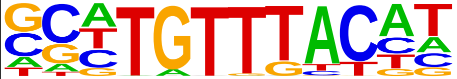 | FOXK2(Forkhead)/U2OS-FOXK2-ChIP-Seq(E-MTAB-2204)/Homer | 1e-4 | -9.500e+00 | 0.0005 | 1210.0 | 32.10% | 83490.1 | 29.25% | motif file (matrix) | svg |
| 64 | 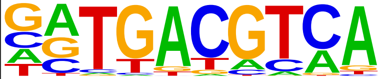 | Atf1(bZIP)/K562-ATF1-ChIP-Seq(GSE31477)/Homer | 1e-3 | -8.046e+00 | 0.0022 | 1111.0 | 29.47% | 76977.4 | 26.97% | motif file (matrix) | svg |
| 65 | 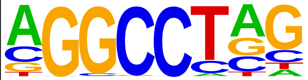 | ZNF711(Zf)/SHSY5Y-ZNF711-ChIP-Seq(GSE20673)/Homer | 1e-3 | -7.506e+00 | 0.0037 | 2156.0 | 57.19% | 155659.5 | 54.53% | motif file (matrix) | svg |
| 66 | 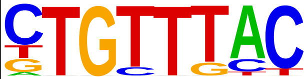 | Foxo1(Forkhead)/RAW-Foxo1-ChIP-Seq(Fan_et_al.)/Homer | 1e-3 | -7.437e+00 | 0.0039 | 2669.0 | 70.80% | 195073.5 | 68.34% | motif file (matrix) | svg |
| 67 | 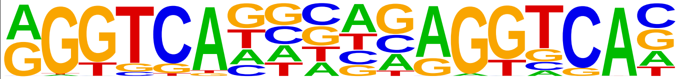 | RAR:RXR(NR),DR5/ES-RAR-ChIP-Seq(GSE56893)/Homer | 1e-3 | -7.169e+00 | 0.0051 | 71.0 | 1.88% | 3591.4 | 1.26% | motif file (matrix) | svg |
| 68 | 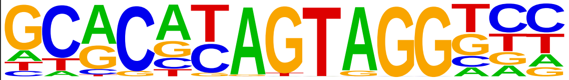 | ZKSCAN1(Zf)/HepG2-ZKSCAN1-ChIP-Seq(Encode)/Homer | 1e-2 | -6.656e+00 | 0.0083 | 80.0 | 2.12% | 4231.9 | 1.48% | motif file (matrix) | svg |
| 69 | 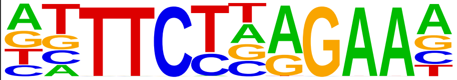 | STAT5(Stat)/mCD4+-Stat5-ChIP-Seq(GSE12346)/Homer | 1e-2 | -6.554e+00 | 0.0091 | 684.0 | 18.14% | 46560.3 | 16.31% | motif file (matrix) | svg |
| 70 | 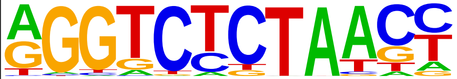 | PRDM14(Zf)/H1-PRDM14-ChIP-Seq(GSE22767)/Homer | 1e-2 | -6.494e+00 | 0.0095 | 618.0 | 16.39% | 41811.6 | 14.65% | motif file (matrix) | svg |
| 71 | 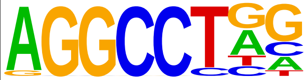 | ZFX(Zf)/mES-Zfx-ChIP-Seq(GSE11431)/Homer | 1e-2 | -6.308e+00 | 0.0113 | 1758.0 | 46.63% | 126345.0 | 44.26% | motif file (matrix) | svg |
| 72 | 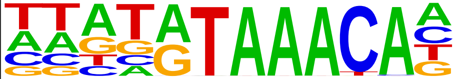 | Foxf1(Forkhead)/Lung-Foxf1-ChIP-Seq(GSE77951)/Homer | 1e-2 | -6.196e+00 | 0.0125 | 1690.0 | 44.83% | 121305.6 | 42.50% | motif file (matrix) | svg |
| 73 | 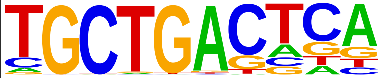 | MafA(bZIP)/Islet-MafA-ChIP-Seq(GSE30298)/Homer | 1e-2 | -5.822e+00 | 0.0178 | 1411.0 | 37.43% | 100658.6 | 35.26% | motif file (matrix) | svg |
| 74 | 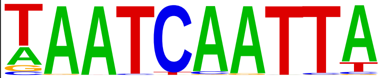 | Pax7(Paired,Homeobox)/Myoblast-Pax7-ChIP-Seq(GSE25064)/Homer | 1e-2 | -5.764e+00 | 0.0187 | 197.0 | 5.23% | 12232.4 | 4.29% | motif file (matrix) | svg |
| 75 | 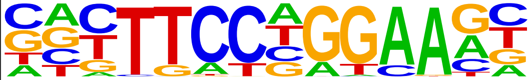 | Stat3+il21(Stat)/CD4-Stat3-ChIP-Seq(GSE19198)/Homer | 1e-2 | -5.724e+00 | 0.0192 | 1251.0 | 33.18% | 88797.5 | 31.11% | motif file (matrix) | svg |
| 76 | 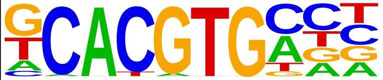 | bHLHE40(bHLH)/HepG2-BHLHE40-ChIP-Seq(GSE31477)/Homer | 1e-2 | -5.638e+00 | 0.0206 | 496.0 | 13.16% | 33434.1 | 11.71% | motif file (matrix) | svg |
| 77 | 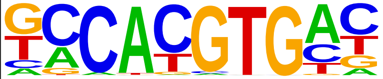 | NPAS2(bHLH)/Liver-NPAS2-ChIP-Seq(GSE39860)/Homer | 1e-2 | -5.630e+00 | 0.0206 | 1753.0 | 46.50% | 126472.9 | 44.31% | motif file (matrix) | svg |
| 78 | 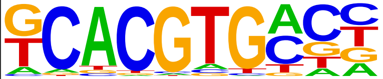 | bHLHE41(bHLH)/proB-Bhlhe41-ChIP-Seq(GSE93764)/Homer | 1e-2 | -5.625e+00 | 0.0206 | 1535.0 | 40.72% | 110086.7 | 38.57% | motif file (matrix) | svg |

| | | | | | | | | | | | |
|---|---|---|---|---|---|---|---|---|---|---|---|
| 79 | 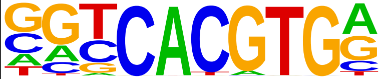 | USF1(bHLH)/GM12878-Usf1-ChIP-Seq(GSE32465)/Homer | 1e-2 | -5.622e+00 | 0.0206 | 747.0 | 19.81% | 51664.7 | 18.10% | motif file (matrix) | svg |
| 80 | 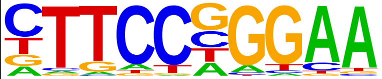 | Stat3(Stat)/mES-Stat3-ChIP-Seq(GSE11431)/Homer | 1e-2 | -4.766e+00 | 0.0468 | 881.0 | 23.37% | 62058.7 | 21.74% | motif file (matrix) | svg |
| 81 | 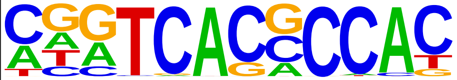 | Srebp2(bHLH)/HepG2-Srebp2-ChIP-Seq(GSE31477)/Homer | 1e-2 | -4.705e+00 | 0.0492 | 238.0 | 6.31% | 15446.8 | 5.41% | motif file (matrix) | svg |
