## Supplementary material for "Reversible and Causal Epigenetic Information Loss in Liver Aging and Disease": Supp. Table 6

| Rank | Motif | Name | P-value | log P-pvalue | q-value (Benjamini) | # Target Sequences with Motif | % of Targets Sequences with Motif | # Background Sequences with Motif | % of Background Sequences with Motif | Motif File | SVG |
|---|---|---|---|---|---|---|---|---|---|---|---|
| 1 | 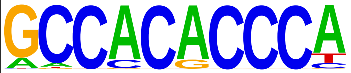 | Klf4(Zf)/mES-Klf4-ChIP-Seq(GSE11431)/Homer | 1e-17 | -4.126e+01 | 0.0000 | 667.0 | 29.45% | 59176.2 | 21.59% | motif file (matrix) | svg |
| 2 | 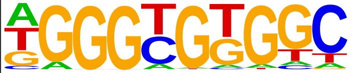 | KLF5(Zf)/LoVo-KLF5-ChIP-Seq(GSE49402)/Homer | 1e-16 | -3.820e+01 | 0.0000 | 1493.0 | 65.92% | 157027.4 | 57.28% | motif file (matrix) | svg |
| 3 | 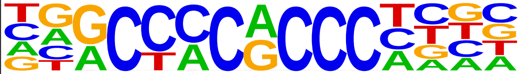 | KLF3(Zf)/MEF-Klf3-ChIP-Seq(GSE44748)/Homer | 1e-16 | -3.728e+01 | 0.0000 | 840.0 | 37.09% | 79473.1 | 28.99% | motif file (matrix) | svg |
| 4 | 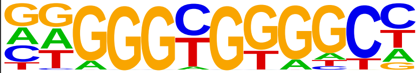 | KLF1(Zf)/HUDEP2-KLF1-CutnRun(GSE136251)/Homer | 1e-16 | -3.685e+01 | 0.0000 | 1209.0 | 53.38% | 122637.8 | 44.73% | motif file (matrix) | svg |
| 5 | 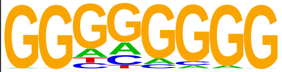 | Maz(Zf)/HepG2-Maz-ChIP-Seq(GSE31477)/Homer | 1e-15 | -3.646e+01 | 0.0000 | 1534.0 | 67.73% | 162786.2 | 59.38% | motif file (matrix) | svg |
| 6 | 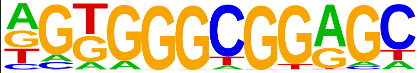 | Sp5(Zf)/mES-Sp5.Flag-ChIP-Seq(GSE72989)/Homer | 1e-14 | -3.366e+01 | 0.0000 | 1295.0 | 57.17% | 134157.8 | 48.94% | motif file (matrix) | svg |
| 7 | 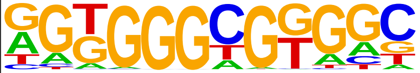 | KLF14(Zf)/HEK293-KLF14.GFP-ChIP-Seq(GSE58341)/Homer | 1e-14 | -3.346e+01 | 0.0000 | 1736.0 | 76.64% | 189825.3 | 69.24% | motif file (matrix) | svg |
| 8 | 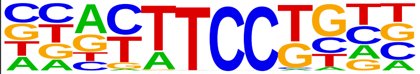 | Etv2(ETS)/ES-ER71-ChIP-Seq(GSE59402)/Homer | 1e-14 | -3.290e+01 | 0.0000 | 1114.0 | 49.18% | 112687.8 | 41.10% | motif file (matrix) | svg |
| 9 | 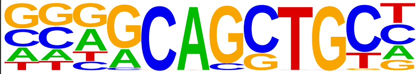 | Ascl2(bHLH)/ESC-Ascl2-ChIP-Seq(GSE97712)/Homer | 1e-13 | -3.011e+01 | 0.0000 | 1289.0 | 56.91% | 134749.4 | 49.15% | motif file (matrix) | svg |
| 10 | 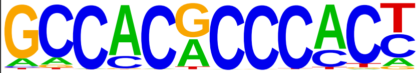 | Klf9(Zf)/GBM-Klf9-ChIP-Seq(GSE62211)/Homer | 1e-12 | -2.913e+01 | 0.0000 | 694.0 | 30.64% | 65642.8 | 23.94% | motif file (matrix) | svg |
| 11 | 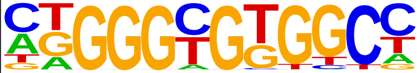 | KLF6(Zf)/PDAC-KLF6-ChIP-Seq(GSE64557)/Homer | 1e-12 | -2.880e+01 | 0.0000 | 1284.0 | 56.69% | 134651.4 | 49.12% | motif file (matrix) | svg |
| 12 | 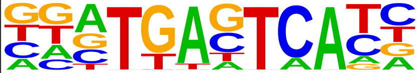 | Fos(bZIP)/TSC-Fos-ChIP-Seq(GSE110950)/Homer | 1e-12 | -2.878e+01 | 0.0000 | 611.0 | 26.98% | 56568.0 | 20.63% | motif file (matrix) | svg |
| 13 | 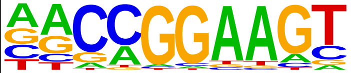 | GABPA(ETS)/Jurkat-GABPa-ChIP-Seq(GSE17954)/Homer | 1e-12 | -2.843e+01 | 0.0000 | 1102.0 | 48.65% | 112925.8 | 41.19% | motif file (matrix) | svg |
| 14 | 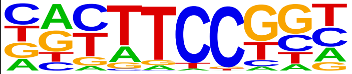 | Fli1(ETS)/CD8-FLI-ChIP-Seq(GSE20898)/Homer | 1e-12 | -2.811e+01 | 0.0000 | 1270.0 | 56.07% | 133222.1 | 48.59% | motif file (matrix) | svg |
| 15 | 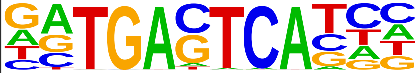 | Jun-AP1(bZIP)/K562-cJun-ChIP-Seq(GSE31477)/Homer | 1e-12 | -2.770e+01 | 0.0000 | 304.0 | 13.42% | 24418.9 | 8.91% | motif file (matrix) | svg |
| 16 | 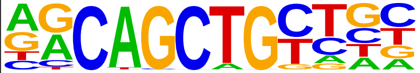 | MyoD(bHLH)/Myotube-MyoD-ChIP-Seq(GSE21614)/Homer | 1e-11 | -2.761e+01 | 0.0000 | 967.0 | 42.69% | 97354.3 | 35.51% | motif file (matrix) | svg |

| | | | | | | | | | | | |
|---|---|---|---|---|---|---|---|---|---|---|---|
| 17 | 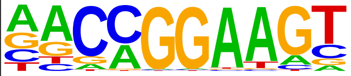 | ETV1(ETS)/GIST48-ETV1-ChIP-Seq(GSE22441)/Homer | 1e-11 | -2.732e+01 | 0.0000 | 1450.0 | 64.02% | 155661.0 | 56.78% | motif file (matrix) | svg |
| 18 | 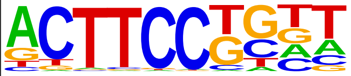 | Elf4(ETS)/BMDM-Elf4-ChIP-Seq(GSE88699)/Homer | 1e-11 | -2.718e+01 | 0.0000 | 1187.0 | 52.41% | 123570.5 | 45.07% | motif file (matrix) | svg |
| 19 | 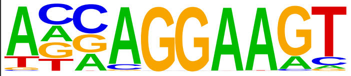 | ELF3(ETS)/PDAC-ELF3-ChIP-Seq(GSE64557)/Homer | 1e-11 | -2.668e+01 | 0.0000 | 891.0 | 39.34% | 88882.3 | 32.42% | motif file (matrix) | svg |
| 20 | 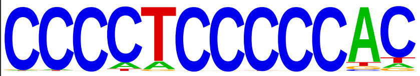 | Zfp281(Zf)/ES-Zfp281-ChIP-Seq(GSE81042)/Homer | 1e-11 | -2.596e+01 | 0.0000 | 520.0 | 22.96% | 47499.8 | 17.33% | motif file (matrix) | svg |
| 21 | 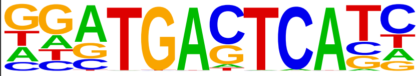 | Fra2(bZIP)/Striatum-Fra2-ChIP-Seq(GSE43429)/Homer | 1e-10 | -2.436e+01 | 0.0000 | 512.0 | 22.60% | 47122.1 | 17.19% | motif file (matrix) | svg |
| 22 | 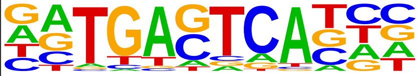 | Fosl2(bZIP)/3T3L1-Fosl2-ChIP-Seq(GSE56872)/Homer | 1e-10 | -2.398e+01 | 0.0000 | 389.0 | 17.17% | 34066.9 | 12.43% | motif file (matrix) | svg |
| 23 | 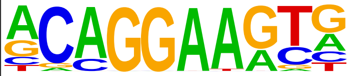 | ERG(ETS)/VCaP-ERG-ChIP-Seq(GSE14097)/Homer | 1e-10 | -2.312e+01 | 0.0000 | 1546.0 | 68.26% | 169436.2 | 61.80% | motif file (matrix) | svg |
| 24 | 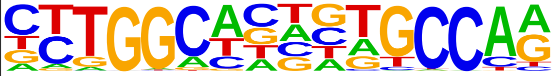 | NF1(CTF)/LNCAP-NF1-ChIP-Seq(Unpublished)/Homer | 1e-10 | -2.307e+01 | 0.0000 | 560.0 | 24.72% | 52794.1 | 19.26% | motif file (matrix) | svg |
| 25 | 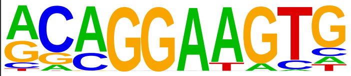 | ETS1(ETS)/Jurkat-ETS1-ChIP-Seq(GSE17954)/Homer | 1e-9 | -2.285e+01 | 0.0000 | 1181.0 | 52.14% | 124669.8 | 45.47% | motif file (matrix) | svg |
| 26 | 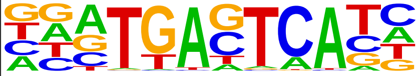 | Fra1(bZIP)/BT549-Fra1-ChIP-Seq(GSE46166)/Homer | 1e-9 | -2.261e+01 | 0.0000 | 565.0 | 24.94% | 53502.5 | 19.52% | motif file (matrix) | svg |
| 27 | 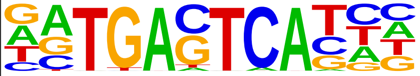 | Atf3(bZIP)/GBM-ATF3-ChIP-Seq(GSE33912)/Homer | 1e-9 | -2.239e+01 | 0.0000 | 652.0 | 28.79% | 63267.7 | 23.08% | motif file (matrix) | svg |
| 28 | 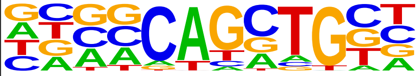 | Ascl1(bHLH)/NeuralTubes-Ascl1-ChIP-Seq(GSE55840)/Homer | 1e-9 | -2.208e+01 | 0.0000 | 1533.0 | 67.68% | 168257.9 | 61.37% | motif file (matrix) | svg |
| 29 | 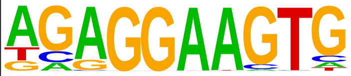 | PU.1(ETS)/ThioMac-PU.1-ChIP-Seq(GSE21512)/Homer | 1e-9 | -2.200e+01 | 0.0000 | 644.0 | 28.43% | 62514.1 | 22.80% | motif file (matrix) | svg |
| 30 | 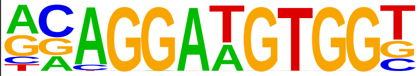 | ETS:RUNX(ETS,Runt)/Jurkat-RUNX1-ChIP-Seq(GSE17954)/Homer | 1e-9 | -2.197e+01 | 0.0000 | 170.0 | 7.51% | 12440.4 | 4.54% | motif file (matrix) | svg |
| 31 | 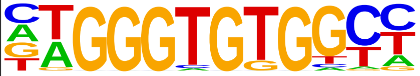 | EKLF(Zf)/Erythrocyte-Klf1-ChIP-Seq(GSE20478)/Homer | 1e-9 | -2.193e+01 | 0.0000 | 365.0 | 16.11% | 32104.3 | 11.71% | motif file (matrix) | svg |
| 32 | 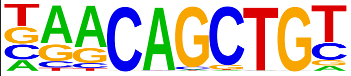 | Myf5(bHLH)/GM-Myf5-ChIP-Seq(GSE24852)/Homer | 1e-9 | -2.186e+01 | 0.0000 | 862.0 | 38.06% | 87437.1 | 31.89% | motif file (matrix) | svg |
| 33 | 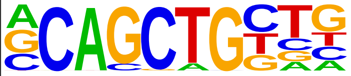 | Tcf12(bHLH)/GM12878-Tcf12-ChIP-Seq(GSE32465)/Homer | 1e-9 | -2.180e+01 | 0.0000 | 1113.0 | 49.14% | 116987.2 | 42.67% | motif file (matrix) | svg |
| 34 | 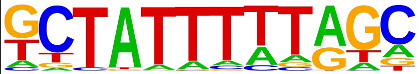 | Mef2d(MADS)/Retina-Mef2d-ChIP-Seq(GSE61391)/Homer | 1e-9 | -2.121e+01 | 0.0000 | 228.0 | 10.07% | 18221.0 | 6.65% | motif file (matrix) | svg |

| | | | | | | | | | | | |
|---|---|---|---|---|---|---|---|---|---|---|---|
| 35 | 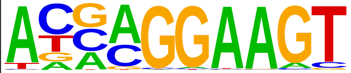 | ELF5(ETS)/T47D-ELF5-ChIP-Seq(GSE30407)/Homer | 1e-9 | -2.117e+01 | 0.0000 | 878.0 | 38.76% | 89584.1 | 32.68% | motif file (matrix) | svg |
| 36 | 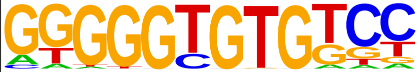 | KLF10(Zf)/HEK293-KLF10.GFP-ChIP-Seq(GSE58341)/Homer | 1e-8 | -2.057e+01 | 0.0000 | 951.0 | 41.99% | 98369.5 | 35.88% | motif file (matrix) | svg |
| 37 | 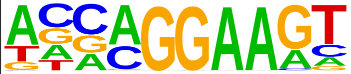 | EHF(ETS)/LoVo-EHF-ChIP-Seq(GSE49402)/Homer | 1e-8 | -2.042e+01 | 0.0000 | 1332.0 | 58.81% | 144111.9 | 52.57% | motif file (matrix) | svg |
| 38 | 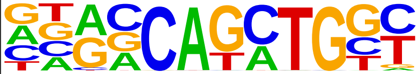 | Atoh1(bHLH)/Cerebellum-Atoh1-ChIP-Seq(GSE22111)/Homer | 1e-8 | -2.024e+01 | 0.0000 | 1199.0 | 52.94% | 128028.8 | 46.70% | motif file (matrix) | svg |
| 39 | 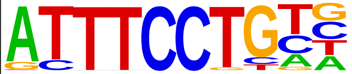 | EWS:ERG-fusion(ETS)/CADO_ES1-EWS:ERG-ChIP-Seq(SRA014231)/Homer | 1e-8 | -1.981e+01 | 0.0000 | 819.0 | 36.16% | 83332.5 | 30.40% | motif file (matrix) | svg |
| 40 | 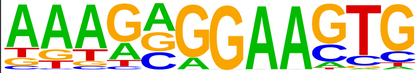 | SpiB(ETS)/OCILY3-SPIB-ChIP-Seq(GSE56857)/Homer | 1e-8 | -1.915e+01 | 0.0000 | 329.0 | 14.53% | 29099.1 | 10.61% | motif file (matrix) | svg |
| 41 | 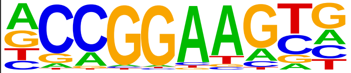 | ETV4(ETS)/HepG2-ETV4-ChIP-Seq(ENCODE)/Homer | 1e-8 | -1.889e+01 | 0.0000 | 1256.0 | 55.45% | 135581.9 | 49.46% | motif file (matrix) | svg |
| 42 | 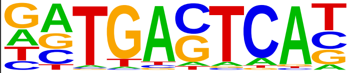 | JunB(bZIP)/DendriticCells-Junb-ChIP-Seq(GSE36099)/Homer | 1e-7 | -1.836e+01 | 0.0000 | 551.0 | 24.33% | 53490.1 | 19.51% | motif file (matrix) | svg |
| 43 | 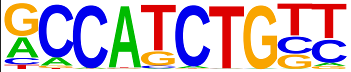 | NeuroD1(bHLH)/Islet-NeuroD1-ChIP-Seq(GSE30298)/Homer | 1e-7 | -1.823e+01 | 0.0000 | 971.0 | 42.87% | 101801.5 | 37.13% | motif file (matrix) | svg |
| 44 | 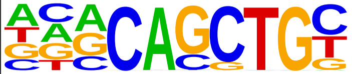 | E2A(bHLH)/proBcell-E2A-ChIP-Seq(GSE21978)/Homer | 1e-7 | -1.759e+01 | 0.0000 | 1465.0 | 64.68% | 161898.4 | 59.05% | motif file (matrix) | svg |
| 45 | 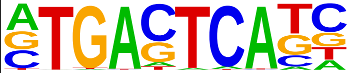 | AP-1(bZIP)/ThioMac-PU.1-ChIP-Seq(GSE21512)/Homer | 1e-7 | -1.710e+01 | 0.0000 | 708.0 | 31.26% | 71763.3 | 26.18% | motif file (matrix) | svg |
| 46 | 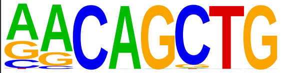 | MyoG(bHLH)/C2C12-MyoG-ChIP-Seq(GSE36024)/Homer | 1e-7 | -1.706e+01 | 0.0000 | 1156.0 | 51.04% | 124419.5 | 45.38% | motif file (matrix) | svg |
| 47 | 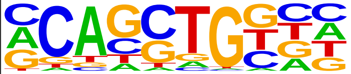 | HEB(bHLH)/mES-Heb-ChIP-Seq(GSE53233)/Homer | 1e-7 | -1.705e+01 | 0.0000 | 1697.0 | 74.92% | 191380.5 | 69.81% | motif file (matrix) | svg |
| 48 | 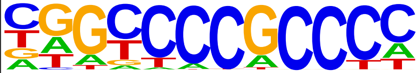 | Sp2(Zf)/HEK293-Sp2.eGFP-ChIP-Seq(Encode)/Homer | 1e-6 | -1.611e+01 | 0.0000 | 1531.0 | 67.59% | 170882.2 | 62.33% | motif file (matrix) | svg |
| 49 | 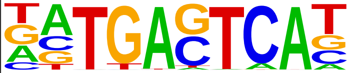 | BATF(bZIP)/Th17-BATF-ChIP-Seq(GSE39756)/Homer | 1e-6 | -1.604e+01 | 0.0000 | 625.0 | 27.59% | 62773.0 | 22.90% | motif file (matrix) | svg |
| 50 | 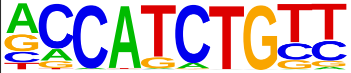 | NeuroG2(bHLH)/Fibroblast-NeuroG2-ChIP-Seq(GSE75910)/Homer | 1e-6 | -1.525e+01 | 0.0000 | 1455.0 | 64.24% | 161888.5 | 59.05% | motif file (matrix) | svg |
| 51 | 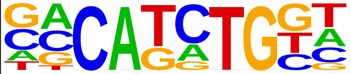 | TCF4(bHLH)/SHSY5Y-TCF4-ChIP-Seq(GSE96915)/Homer | 1e-6 | -1.509e+01 | 0.0000 | 1451.0 | 64.06% | 161482.3 | 58.90% | motif file (matrix) | svg |
| 52 | 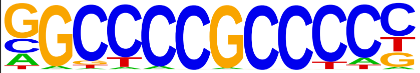 | Sp1(Zf)/Promoter/Homer | 1e-6 | -1.483e+01 | 0.0000 | 446.0 | 19.69% | 43214.0 | 15.76% | motif file (matrix) | svg |

| | | | | | | | | | | | |
|---|---|---|---|---|---|---|---|---|---|---|---|
| 53 | 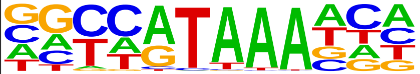 | CDX4(Homeobox)/ZebrafishEmbryos-Cdx4.Myc-ChIP-Seq(GSE48254)/Homer | 1e-6 | -1.437e+01 | 0.0000 | 815.0 | 35.98% | 85447.5 | 31.17% | motif file (matrix) | svg |
| 54 | 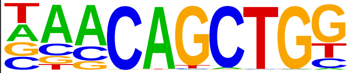 | Tcf21(bHLH)/ArterySmoothMuscle-Tcf21-ChIP-Seq(GSE61369)/Homer | 1e-6 | -1.416e+01 | 0.0000 | 1065.0 | 47.02% | 115074.7 | 41.97% | motif file (matrix) | svg |
| 55 | 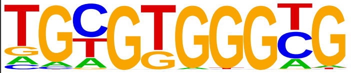 | Egr1(Zf)/K562-Egr1-ChIP-Seq(GSE32465)/Homer | 1e-6 | -1.414e+01 | 0.0000 | 888.0 | 39.21% | 94113.7 | 34.33% | motif file (matrix) | svg |
| 56 | 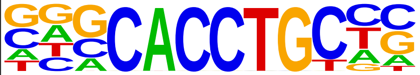 | Slug(Zf)/Mesoderm-Snai2-ChIP-Seq(GSE61475)/Homer | 1e-5 | -1.300e+01 | 0.0000 | 725.0 | 32.01% | 75710.2 | 27.62% | motif file (matrix) | svg |
| 57 | 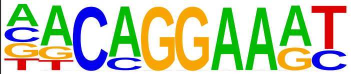 | EWS:FLI1-fusion(ETS)/SK_N_MC-EWS:FLI1-ChIP-Seq(SRA014231)/Homer | 1e-5 | -1.300e+01 | 0.0000 | 716.0 | 31.61% | 74670.9 | 27.24% | motif file (matrix) | svg |
| 58 | 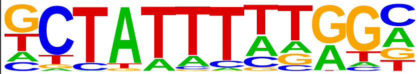 | Mef2b(MADS)/HEK293-Mef2b.V5-ChIP-Seq(GSE67450)/Homer | 1e-5 | -1.273e+01 | 0.0000 | 736.0 | 32.49% | 77130.5 | 28.13% | motif file (matrix) | svg |
| 59 | 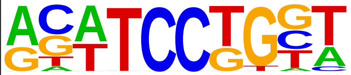 | SPDEF(ETS)/VCaP-SPDEF-ChIP-Seq(SRA014231)/Homer | 1e-5 | -1.264e+01 | 0.0000 | 1114.0 | 49.18% | 121832.2 | 44.44% | motif file (matrix) | svg |
| 60 | 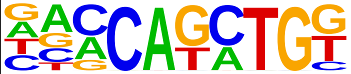 | BHLHA15(bHLH)/NIH3T3-BHLHB8.HA-ChIP-Seq(GSE119782)/Homer | 1e-5 | -1.237e+01 | 0.0000 | 1362.0 | 60.13% | 152098.6 | 55.48% | motif file (matrix) | svg |
| 61 | 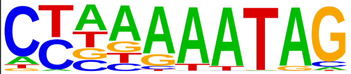 | Mef2a(MADS)/HL1-Mef2a.biotin-ChIP-Seq(GSE21529)/Homer | 1e-5 | -1.173e+01 | 0.0001 | 397.0 | 17.53% | 39062.7 | 14.25% | motif file (matrix) | svg |
| 62 | 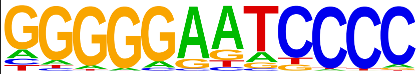 | NFkB-p50,p52(RHD)/Monocyte-p50-ChIP-Chip(Schreiber_et_al.)/Homer | 1e-5 | -1.172e+01 | 0.0001 | 171.0 | 7.55% | 14735.5 | 5.37% | motif file (matrix) | svg |
| 63 | 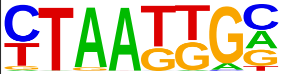 | Isl1(Homeobox)/Neuron-Isl1-ChIP-Seq(GSE31456)/Homer | 1e-5 | -1.169e+01 | 0.0001 | 1493.0 | 65.92% | 168672.1 | 61.53% | motif file (matrix) | svg |
| 64 | 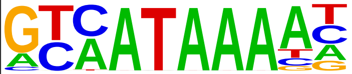 | Cdx2(Homeobox)/mES-Cdx2-ChIP-Seq(GSE14586)/Homer | 1e-4 | -1.148e+01 | 0.0001 | 647.0 | 28.57% | 67516.5 | 24.63% | motif file (matrix) | svg |
| 65 | 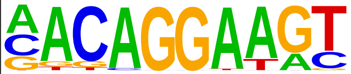 | Ets1-distal(ETS)/CD4+-PolII-ChIP-Seq(Barski_et_al.)/Homer | 1e-4 | -1.142e+01 | 0.0001 | 409.0 | 18.06% | 40534.8 | 14.79% | motif file (matrix) | svg |
| 66 | 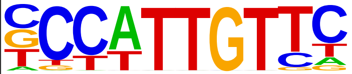 | Sox2(HMG)/mES-Sox2-ChIP-Seq(GSE11431)/Homer | 1e-4 | -1.121e+01 | 0.0001 | 859.0 | 37.92% | 92385.0 | 33.70% | motif file (matrix) | svg |
| 67 | 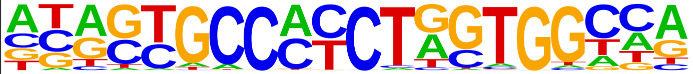 | CTCF(Zf)/CD4+-CTCF-ChIP-Seq(Barski_et_al.)/Homer | 1e-4 | -1.109e+01 | 0.0001 | 251.0 | 11.08% | 23350.6 | 8.52% | motif file (matrix) | svg |
| 68 | 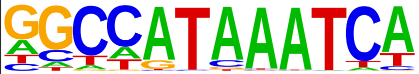 | Hoxc9(Homeobox)/Ainv15-Hoxc9-ChIP-Seq(GSE21812)/Homer | 1e-4 | -1.054e+01 | 0.0002 | 490.0 | 21.63% | 50063.2 | 18.26% | motif file (matrix) | svg |
| 69 | 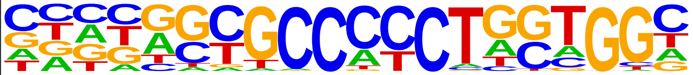 | BORIS(Zf)/K562-CTCFL-ChIP-Seq(GSE32465)/Homer | 1e-4 | -1.029e+01 | 0.0002 | 353.0 | 15.58% | 34810.7 | 12.70% | motif file (matrix) | svg |
| 70 | 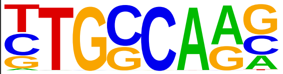 | NF1-halfsite(CTF)/LNCaP-NF1-ChIP-Seq(Unpublished)/Homer | 1e-4 | -1.002e+01 | 0.0003 | 1547.0 | 68.30% | 176464.9 | 64.37% | motif file (matrix) | svg |

| | | | | | | | | | | | |
|---|---|---|---|---|---|---|---|---|---|---|---|
| 71 | 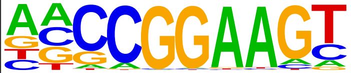 | ELF1(ETS)/Jurkat-ELF1-ChIP-Seq(SRA014231)/Homer | 1e-4 | -9.893e+00 | 0.0003 | 676.0 | 29.85% | 71785.2 | 26.18% | motif file (matrix) | svg |
| 72 | 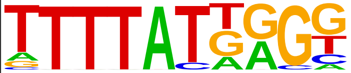 | HOXB13(Homeobox)/ProstateTumor-HOXB13-ChIP-Seq(GSE56288)/Homer | 1e-4 | -9.882e+00 | 0.0003 | 930.0 | 41.06% | 101627.2 | 37.07% | motif file (matrix) | svg |
| 73 | 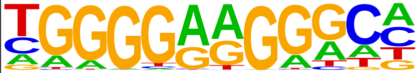 | ZNF467(Zf)/HEK293-ZNF467.GFP-ChIP-Seq(GSE58341)/Homer | 1e-4 | -9.763e+00 | 0.0003 | 1087.0 | 47.99% | 120454.8 | 43.94% | motif file (matrix) | svg |
| 74 | 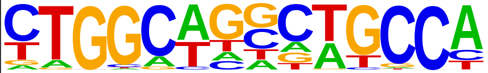 | Tlx?(NR)/NPC-H3K4me1-ChIP-Seq(GSE16256)/Homer | 1e-4 | -9.598e+00 | 0.0004 | 559.0 | 24.68% | 58446.7 | 21.32% | motif file (matrix) | svg |
| 75 | 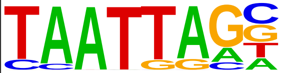 | Lhx2(Homeobox)/HFSC-Lhx2-ChIP-Seq(GSE48068)/Homer | 1e-4 | -9.470e+00 | 0.0005 | 900.0 | 39.74% | 98348.2 | 35.87% | motif file (matrix) | svg |
| 76 | 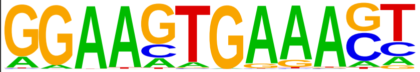 | PU.1:IRF8(ETS:IRF)/pDC-Irf8-ChIP-Seq(GSE66899)/Homer | 1e-3 | -9.155e+00 | 0.0006 | 239.0 | 10.55% | 22771.7 | 8.31% | motif file (matrix) | svg |
| 77 | 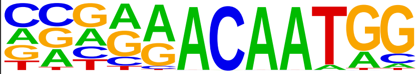 | Sox7(HMG)/ESC-Sox7-ChIP-Seq(GSE133899)/Homer | 1e-3 | -8.734e+00 | 0.0009 | 330.0 | 14.57% | 32966.3 | 12.02% | motif file (matrix) | svg |
| 78 | 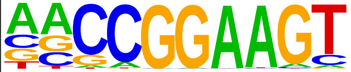 | ETS(ETS)/Promoter/Homer | 1e-3 | -8.678e+00 | 0.0010 | 453.0 | 20.00% | 46850.5 | 17.09% | motif file (matrix) | svg |
| 79 | 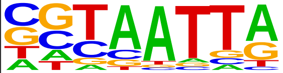 | DLX5(Homeobox)/BasalGanglia-Dlx5-ChIP-seq(GSE124936)/Homer | 1e-3 | -8.597e+00 | 0.0010 | 686.0 | 30.29% | 73770.2 | 26.91% | motif file (matrix) | svg |
| 80 | 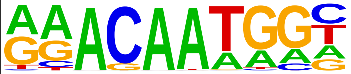 | Sox15(HMG)/CPA-Sox15-ChIP-Seq(GSE62909)/Homer | 1e-3 | -8.493e+00 | 0.0011 | 980.0 | 43.27% | 108557.5 | 39.60% | motif file (matrix) | svg |
| 81 | 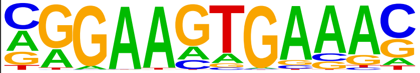 | PU.1-IRF(ETS:IRF)/Bcell-PU.1-ChIP-Seq(GSE21512)/Homer | 1e-3 | -8.428e+00 | 0.0012 | 1269.0 | 56.03% | 143434.1 | 52.32% | motif file (matrix) | svg |
| 82 | 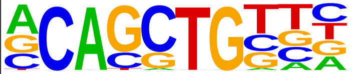 | Ptf1a(bHLH)/Panc1-Ptf1a-ChIP-Seq(GSE47459)/Homer | 1e-3 | -8.165e+00 | 0.0015 | 1916.0 | 84.59% | 224338.5 | 81.83% | motif file (matrix) | svg |
| 83 | 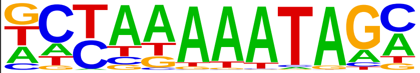 | Mef2c(MADS)/GM12878-Mef2c-ChIP-Seq(GSE32465)/Homer | 1e-3 | -8.154e+00 | 0.0015 | 408.0 | 18.01% | 42030.9 | 15.33% | motif file (matrix) | svg |
| 84 | 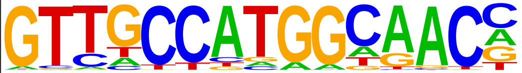 | Rfx2(HTH)/LoVo-RFX2-ChIP-Seq(GSE49402)/Homer | 1e-3 | -7.930e+00 | 0.0019 | 153.0 | 6.75% | 13989.5 | 5.10% | motif file (matrix) | svg |
| 85 | 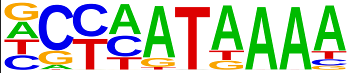 | Hoxd13(Homeobox)/ChickenMSG-Hoxd13.Flag-ChIP-Seq(GSE86088)/Homer | 1e-3 | -7.772e+00 | 0.0022 | 1222.0 | 53.95% | 138238.5 | 50.42% | motif file (matrix) | svg |
| 86 | 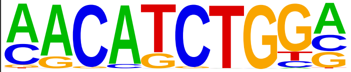 | ZBTB18(Zf)/HEK293-ZBTB18.GFP-ChIP-Seq(GSE58341)/Homer | 1e-3 | -7.771e+00 | 0.0022 | 597.0 | 26.36% | 63961.0 | 23.33% | motif file (matrix) | svg |
| 87 | 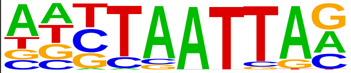 | Lhx3(Homeobox)/Neuron-Lhx3-ChIP-Seq(GSE31456)/Homer | 1e-3 | -7.750e+00 | 0.0022 | 1227.0 | 54.17% | 138862.4 | 50.65% | motif file (matrix) | svg |
| 88 | 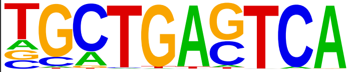 | Bach2(bZIP)/OCILy7-Bach2-ChIP-Seq(GSE44420)/Homer | 1e-3 | -7.729e+00 | 0.0022 | 217.0 | 9.58% | 20948.7 | 7.64% | motif file (matrix) | svg |

| | | | | | | | | | | | |
|---|---|---|---|---|---|---|---|---|---|---|---|
| 89 | 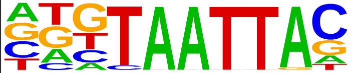 | Dlx3(Homeobox)/Kerainocytes-Dlx3-ChIP-Seq(GSE89884)/Homer | 1e-3 | -7.613e+00 | 0.0024 | 592.0 | 26.14% | 63487.8 | 23.16% | motif file (matrix) | svg |
| 90 | 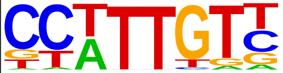 | Sox3(HMG)/NPC-Sox3-ChIP-Seq(GSE33059)/Homer | 1e-3 | -7.523e+00 | 0.0026 | 1393.0 | 61.50% | 159290.3 | 58.10% | motif file (matrix) | svg |
| 91 | 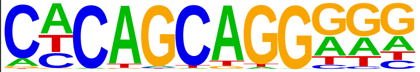 | Unknown-ESC-element(?)/mES-Nanog-ChIP-Seq(GSE11724)/Homer | 1e-3 | -7.499e+00 | 0.0027 | 717.0 | 31.66% | 78161.4 | 28.51% | motif file (matrix) | svg |
| 92 | 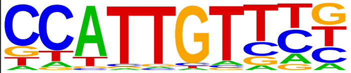 | Sox17(HMG)/Endoderm-Sox17-ChIP-Seq(GSE61475)/Homer | 1e-3 | -7.325e+00 | 0.0032 | 680.0 | 30.02% | 73950.8 | 26.97% | motif file (matrix) | svg |
| 93 | 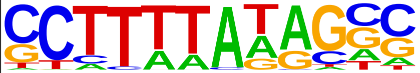 | TATA-Box(TBP)/Promoter/Homer | 1e-3 | -7.293e+00 | 0.0032 | 1160.0 | 51.21% | 131122.3 | 47.83% | motif file (matrix) | svg |
| 94 | 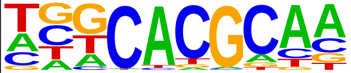 | Arnt:Ahr(bHLH)/MCF7-Arnt-ChIP-Seq(Lo_et_al.)/Homer | 1e-3 | -7.262e+00 | 0.0033 | 707.0 | 31.21% | 77161.4 | 28.15% | motif file (matrix) | svg |
| 95 | 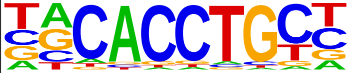 | Snail1(Zf)/LS174T-SNAIL1.HA-ChIP-Seq(GSE127183)/Homer | 1e-3 | -7.175e+00 | 0.0035 | 961.0 | 42.43% | 107311.4 | 39.14% | motif file (matrix) | svg |
| 96 | 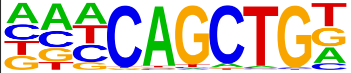 | Ap4(bHLH)/AML-Tfap4-ChIP-Seq(GSE45738)/Homer | 1e-3 | -7.123e+00 | 0.0037 | 1214.0 | 53.60% | 137800.5 | 50.26% | motif file (matrix) | svg |
| 97 | 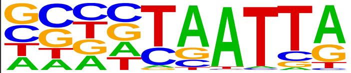 | DLX1(Homeobox)/BasalGanglia-Dlx1-ChIP-seq(GSE124936)/Homer | 1e-3 | -7.008e+00 | 0.0041 | 1048.0 | 46.27% | 117868.5 | 42.99% | motif file (matrix) | svg |
| 98 | 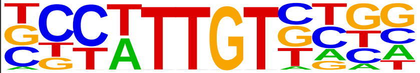 | Sox21(HMG)/ESC-SOX21-ChIP-Seq(GSE110505)/Homer | 1e-2 | -6.887e+00 | 0.0046 | 1457.0 | 64.33% | 167674.9 | 61.16% | motif file (matrix) | svg |
| 99 | 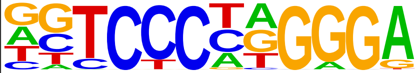 | EBF(EBF)/proBcell-EBF-ChIP-Seq(GSE21978)/Homer | 1e-2 | -6.654e+00 | 0.0057 | 299.0 | 13.20% | 30544.8 | 11.14% | motif file (matrix) | svg |
| 100 | 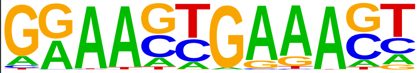 | IRF8(IRF)/BMDM-IRF8-ChIP-Seq(GSE77884)/Homer | 1e-2 | -6.597e+00 | 0.0060 | 359.0 | 15.85% | 37356.8 | 13.63% | motif file (matrix) | svg |
| 101 | 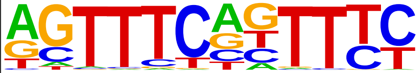 | IRF3(IRF)/BMDM-Irf3-ChIP-Seq(GSE67343)/Homer | 1e-2 | -6.583e+00 | 0.0060 | 349.0 | 15.41% | 36229.2 | 13.22% | motif file (matrix) | svg |
| 102 | 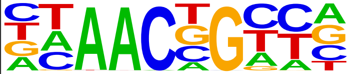 | BMYB(HTH)/Hela-BMYB-ChIP-Seq(GSE27030)/Homer | 1e-2 | -6.560e+00 | 0.0061 | 1314.0 | 58.01% | 150443.8 | 54.88% | motif file (matrix) | svg |
| 103 | 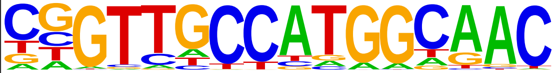 | RFX(HTH)/K562-RFX3-ChIP-Seq(SRA012198)/Homer | 1e-2 | -6.544e+00 | 0.0061 | 137.0 | 6.05% | 12765.9 | 4.66% | motif file (matrix) | svg |
| 104 | 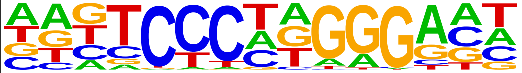 | EBF2(EBF)/BrownAdipose-EBF2-ChIP-Seq(GSE97114)/Homer | 1e-2 | -6.484e+00 | 0.0065 | 1019.0 | 44.99% | 114835.3 | 41.89% | motif file (matrix) | svg |
| 105 | 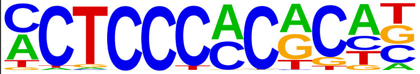 | WT1(Zf)/Kidney-WT1-ChIP-Seq(GSE90016)/Homer | 1e-2 | -6.413e+00 | 0.0069 | 770.0 | 34.00% | 85243.1 | 31.09% | motif file (matrix) | svg |
| 106 | 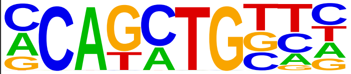 | Twist2(bHLH)/Myoblast-Twist2.Ty1-ChIP-Seq(GSE127998)/Homer | 1e-2 | -6.230e+00 | 0.0082 | 1528.0 | 67.46% | 176986.7 | 64.56% | motif file (matrix) | svg |

| | | | | | | | | | | | |
|---|---|---|---|---|---|---|---|---|---|---|---|
| 107 | 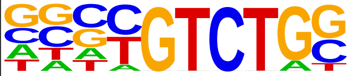 | Smad4(MAD)/ESC-SMAD4-ChIP-Seq(GSE29422)/Homer | 1e-2 | -6.157e+00 | 0.0087 | 1503.0 | 66.36% | 173967.9 | 63.46% | motif file (matrix) | svg |
| 108 | 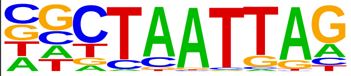 | LHX9(Homeobox)/Hct116-LHX9.V5-ChIP-Seq(GSE116822)/Homer | 1e-2 | -6.025e+00 | 0.0099 | 1124.0 | 49.62% | 127881.8 | 46.65% | motif file (matrix) | svg |
| 109 | 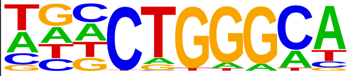 | ZNF416(Zf)/HEK293-ZNF416.GFP-ChIP-Seq(GSE58341)/Homer | 1e-2 | -6.008e+00 | 0.0099 | 1337.0 | 59.03% | 153746.0 | 56.08% | motif file (matrix) | svg |
| 110 | 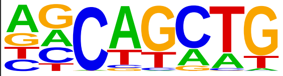 | SCL(bHLH)/HPC7-Scl-ChIP-Seq(GSE13511)/Homer | 1e-2 | -5.898e+00 | 0.0110 | 2193.0 | 96.82% | 262243.5 | 95.66% | motif file (matrix) | svg |
| 111 | 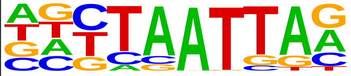 | Lhx1(Homeobox)/EmbryoCarcinoma-Lhx1-ChIP-Seq(GSE70957)/Homer | 1e-2 | -5.767e+00 | 0.0124 | 898.0 | 39.65% | 101002.9 | 36.84% | motif file (matrix) | svg |
| 112 | 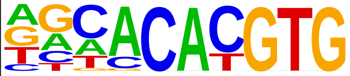 | MNT(bHLH)/HepG2-MNT-ChIP-Seq(Encode)/Homer | 1e-2 | -5.533e+00 | 0.0155 | 982.0 | 43.36% | 111264.6 | 40.59% | motif file (matrix) | svg |
| 113 | 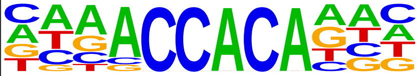 | RUNX2(Runt)/PCa-RUNX2-ChIP-Seq(GSE33889)/Homer | 1e-2 | -5.531e+00 | 0.0155 | 858.0 | 37.88% | 96451.8 | 35.18% | motif file (matrix) | svg |
| 114 | 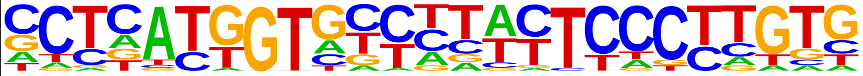 | ZNF41(Zf)/HEK293-ZNF41.GFP-ChIP-Seq(GSE58341)/Homer | 1e-2 | -5.406e+00 | 0.0173 | 39.0 | 1.72% | 2989.1 | 1.09% | motif file (matrix) | svg |
| 115 | 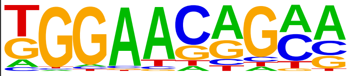 | ZNF189(Zf)/HEK293-ZNF189.GFP-ChIP-Seq(GSE58341)/Homer | 1e-2 | -5.303e+00 | 0.0190 | 906.0 | 40.00% | 102390.6 | 37.35% | motif file (matrix) | svg |
| 116 | 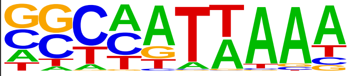 | Unknown(Homeobox)/Limb-p300-ChIP-Seq/Homer | 1e-2 | -5.302e+00 | 0.0190 | 587.0 | 25.92% | 64629.0 | 23.57% | motif file (matrix) | svg |
| 117 | 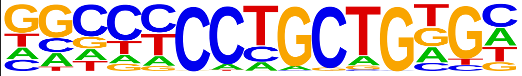 | Zic3(Zf)/mES-Zic3-ChIP-Seq(GSE37889)/Homer | 1e-2 | -5.197e+00 | 0.0208 | 721.0 | 31.83% | 80499.4 | 29.36% | motif file (matrix) | svg |
| 118 | 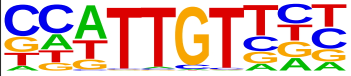 | Sox6(HMG)/Myotubes-Sox6-ChIP-Seq(GSE32627)/Homer | 1e-2 | -5.142e+00 | 0.0218 | 1271.0 | 56.11% | 146543.2 | 53.45% | motif file (matrix) | svg |
| 119 | 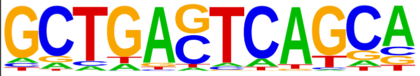 | MafK(bZIP)/C2C12-MafK-ChIP-Seq(GSE36030)/Homer | 1e-2 | -5.100e+00 | 0.0225 | 273.0 | 12.05% | 28494.6 | 10.39% | motif file (matrix) | svg |
| 120 | 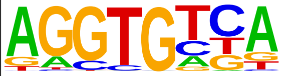 | Tbx5(T-box)/HL1-Tbx5.biotin-ChIP-Seq(GSE21529)/Homer | 1e-2 | -5.081e+00 | 0.0228 | 2010.0 | 88.74% | 238460.7 | 86.98% | motif file (matrix) | svg |
| 121 | 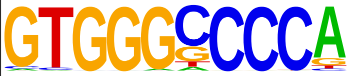 | ZNF692(Zf)/HEK293-ZNF692.GFP-ChIP-Seq(GSE58341)/Homer | 1e-2 | -5.011e+00 | 0.0242 | 186.0 | 8.21% | 18766.1 | 6.85% | motif file (matrix) | svg |
| 122 | 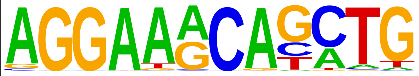 | ETS:E-box(ETS,bHLH)/HPC7-Scl-ChIP-Seq(GSE22178)/Homer | 1e-2 | -4.964e+00 | 0.0252 | 123.0 | 5.43% | 11858.0 | 4.33% | motif file (matrix) | svg |
| 123 | 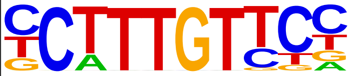 | Sox4(HMG)/proB-Sox4-ChIP-Seq(GSE50066)/Homer | 1e-2 | -4.940e+00 | 0.0256 | 816.0 | 36.03% | 92012.6 | 33.56% | motif file (matrix) | svg |
| 124 | 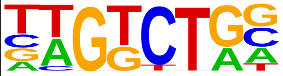 | Smad3(MAD)/NPC-Smad3-ChIP-Seq(GSE36673)/Homer | 1e-2 | -4.891e+00 | 0.0267 | 1973.0 | 87.11% | 233859.2 | 85.30% | motif file (matrix) | svg |

| | | | | | | | | | | | |
|---|---|---|---|---|---|---|---|---|---|---|---|
| 125 | 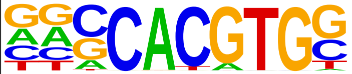 | c-Myc(bHLH)/mES-cMyc-ChIP-Seq(GSE11431)/Homer | 1e-2 | -4.719e+00 | 0.0314 | 564.0 | 24.90% | 62433.3 | 22.77% | motif file (matrix) | svg |
| 126 | 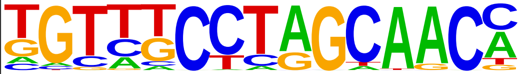 | Rfx6(HTH)/Min6b1-Rfx6.HA-ChIP-Seq(GSE62844)/Homer | 1e-2 | -4.694e+00 | 0.0320 | 1124.0 | 49.62% | 129197.1 | 47.13% | motif file (matrix) | svg |
| 127 | 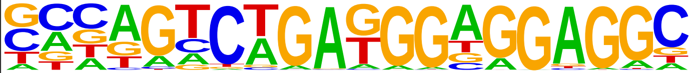 | ZSCAN22(Zf)/HEK293-ZSCAN22.GFP-ChIP-Seq(GSE58341)/Homer | 1e-2 | -4.666e+00 | 0.0326 | 118.0 | 5.21% | 11436.6 | 4.17% | motif file (matrix) | svg |
